# Ibudilast Protects Retinal Bipolar Cells from Excitotoxic Retinal Damage and Activates the mTOR Pathway

**DOI:** 10.1101/2024.03.18.585556

**Authors:** Sumaya Hamadmad, Tyler Heisler-Taylor, Sandeep Goswami, Evan Hawthorn, Sameer Chaurasia, Dena Martini, Diana Summitt, Ali Zaatari, Elizabeth G Urbanski, Kayla Bernstein, Julie Racine, Abhay Satoskar, Heithem M. El-Hodiri, Andy J. Fischer, Colleen M. Cebulla

## Abstract

Ibudilast, an inhibitor of macrophage migration inhibitory factor (MIF) and phosphodiesterase (PDE), has been recently shown to have neuroprotective effects in a variety of neurologic diseases. We utilize a chick excitotoxic retinal damage model to investigate ibudilast’s potential to protect retinal neurons. Using single cell RNA-sequencing (scRNA-seq), we find that MIF, putative MIF receptors CD74 and CD44, and several PDEs are upregulated in different retinal cells during damage. Intravitreal ibudilast is well tolerated in the eye and causes no evidence of toxicity. Ibudilast effectively protects neurons in the inner nuclear layer from NMDA-induced cell death, restores retinal layer thickness on spectral domain optical coherence tomography, and preserves retinal neuron function, particularly for the ON bipolar cells, as assessed by electroretinography. PDE inhibition seems essential for ibudilast’s neuroprotection, as AV1013, the analogue that lacks PDE inhibitor activity, is ineffective. scRNA-seq analysis reveals upregulation of multiple signaling pathways, including mTOR, in damaged Müller glia (MG) with ibudilast treatment compared to AV1013. Components of mTORC1 and mTORC2 are upregulated in both bipolar cells and MG with ibudilast. The mTOR inhibitor rapamycin blocked accumulation of pS6 but did not reduce TUNEL positive dying cells. Additionally, through ligand-receptor interaction analysis, crosstalk between bipolar cells and MG may be important for neuroprotection. We have identified several paracrine signaling pathways that are known to contribute to cell survival and neuroprotection and might play essential roles in ibudilast function. These findings highlight ibudilast’s potential to protect inner retinal neurons during damage and show promise for future clinical translation.

**Graphical Abstract:** *Main Points:*

- Ibudilast, a MIF and PDE inhibitor, preserves the form and function of the retina, especially bipolar cells, during excitotoxic damage
- Ibudilast upregulates multiple signaling pathways, including mTOR, in damaged Müller glia and bipolar cells 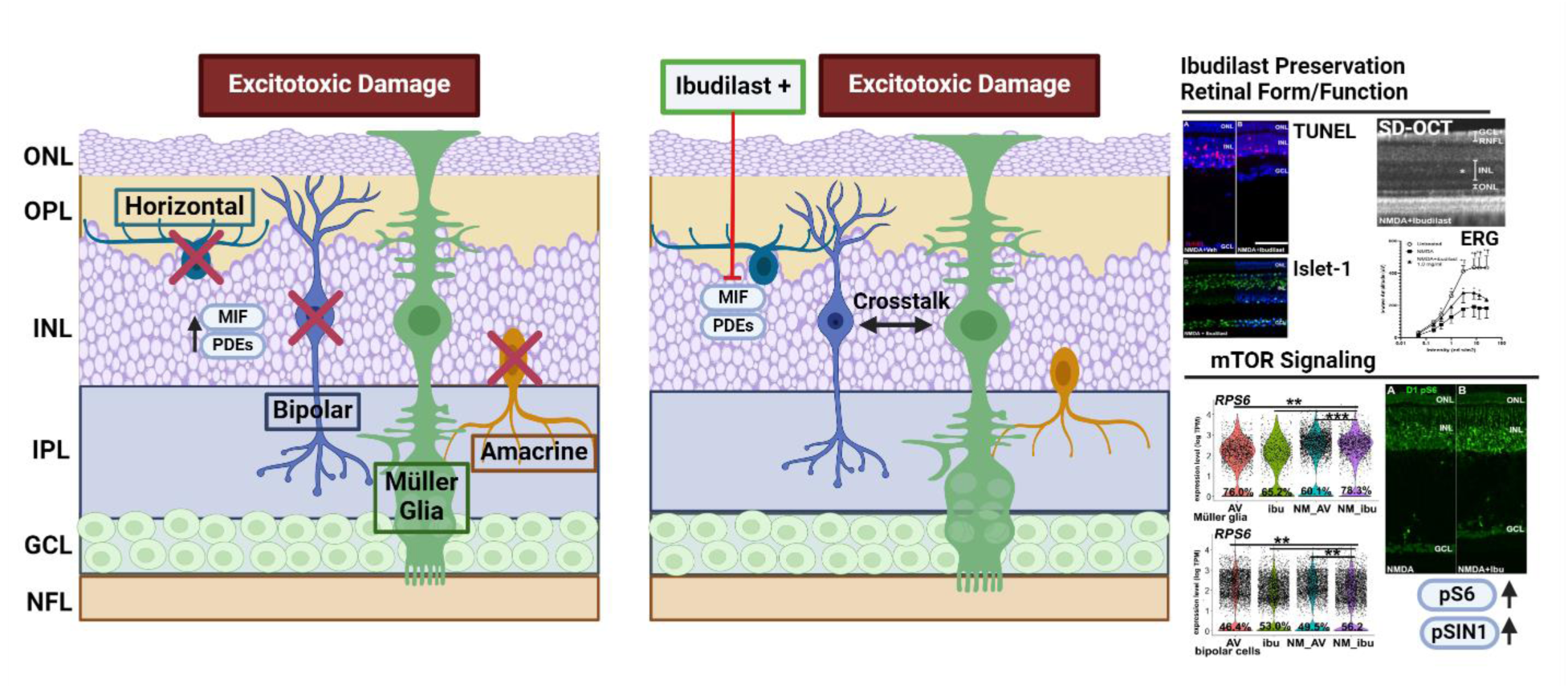

## Introduction

Permanent loss of retinal neurons underlies the vision loss experienced by millions worldwide from pathologic conditions affecting the retina (Flaxman et al., 2017). Retinal maladies that lead to neurodegeneration include mechanisms as diverse as retinal vein occlusion, diabetic retinopathy, retinal detachment, inherited retinal disease, inflammatory conditions, traumatic injury, and glaucoma. A common underlying mechanism of neuronal cell death in these disease and injury states relates to glutamate-mediated excitotoxicity (Ishikawa, 2013; Wisely et al., 2017) where excess glutamate becomes toxic to retinal neurons (Kuehn et al., 2017). Overstimulation of glutamate receptors, including the N-Methyl-D-aspartic acid (NMDA) receptor, leads to uncontrolled calcium ion influx which activates cell death pathways (Smith, 2002). Secondary retinal damage may occur from inflammation, with accumulation of activated microglia and infiltrating macrophages (Fischer, Zelinka, & Milani-Nejad, 2015a; Kuehn et al., 2017). The Müller glia (MG) are also activated by glutamate excitotoxicity and their reactivity can lead to both protective and detrimental effects on retinal neurons. Under normal conditions, MG remove extracellular glutamate from inner retinal tissue through the activity of glutamate uptake carrier GLAST. However, during glutamate toxicity the uptake mechanism can be overwhelmed and reactive MG can respond by undergoing gliosis, upregulating inflammatory factors, recruiting more microglia/macrophages, and further disrupting the glial-neuronal interaction (Bringmann & Wiedemann, 2012). The neuronal-MG interactions during excitotoxicity are not fully understood and are the subject of ongoing research.

We and others have identified the pro-inflammatory cytokine macrophage migration inhibitory factor (MIF) as a mediator of neuronal toxicity in several disease models (Alexander et al., 2012; Kim et al., 2017b)]. MIF plays a key role in immune system regulation and is known to upregulate pro-inflammatory cytokines (David, 1966; Hermanowski-Vosatka et al., 1999; Hudson et al., 1999; Kleemann et al., 2000; O’Reilly, Doroudian, Mawhinney, & Donnelly, 2016; Roger, David, Glauser, & Calandra, 2001; Taguchi, Sugita, Tagawa, Nishihira, & Mochizuki, 2001; Thiele et al., 2015). We have shown that MIF inhibition is anti-gliotic and neuroprotective to photoreceptors in a murine retinal detachment (RD) model (Kim et al., 2017a). Moreover, our lab identified a human genetic association of MIF promoter polymorphisms with epiretinal membrane (ERM) formation (Cebulla et al., 2019). MG are the predominant components of ERM, suggesting that MIF could play an important role in the pathological function of retinal glia (Matsuda, Tagawa, Yoshida, Matsuda, & Nishihira, 1997).

In a search for clinically relevant MIF inhibitors, we identified ibudilast as a drug of interest. Ibudilast is a MIF and phosphodiesterase (PDE) inhibitor, with preferential inhibition of PDE3A, PDE4, PDE10A, and PDE11A (Gibson et al., 2006). Ibudilast is currently approved in Japan for the treatment of post-stroke dizziness and asthma (Cho et al., 2010; Rolan, Hutchinson, & Johnson, 2009). Recently, ibudilast was found to have neuroprotective effects in a variety of neurological disorders (Egashira et al., 2021; Fujita et al., 2018; Zhang et al., 2022). It is in clinical trials in the United States for evaluation as a neuroprotective agent in amyotrophic lateral sclerosis (Oskarsson et al., 2021) and multiple sclerosis (Bermel et al., 2021; Fox et al., 2018; MediciNova, Health, Disorders, Stroke, & Society, 2013; MediciNova & System, 2014; Oskarsson et al., 2021) as well as in drug addiction (Grodin et al., 2022) and pain (Egashira et al., 2021).

PDEs regulate a wide variety of cellular functions by affecting cyclic nucleotide-mediated signaling, including inflammatory responses, cell survival and apoptosis, and synaptic function. Increasing evidence suggest that PDEs are implicated in the pathophysiology of many neuronal diseases (Angelopoulou, Pyrgelis, & Piperi, 2022). Pharmacological screening of several PDE inhibitors showed that PDE4 and PDE5 inhibition significantly relieved mechanical hypersensitivity in a mouse model of peripheral nerve injury (Megat et al., 2022). Additionally, both MIF and PDE4 are involved in neuroinflammatory processes through the regulation of inflammatory responses in microglia (Hertz et al., 2009).

Ibudilast functions as a glial cell modulator and inhibits many other pro-inflammatory molecules, including IL-1, TNFa, IL-6, and toll-like receptor 4 (TLR4) (Angelopoulou et al., 2022), that are upregulated in damaged retinas (Palazzo, Kelly, Koenig, & Fischer, 2023; Todd et al., 2019). The fact that ibudilast can target the neuroinflammatory process by inhibiting both MIF and PDEs makes it an attractive candidate to use as a neuroprotective agent in damaging retinal conditions. Herein we investigate the neuroprotective impact of ibudilast in excitotoxic retinal damage and the underlying signaling pathways impacted by ibudilast and its analog AV1013.

The chick eye disease model has become increasingly popular in recent years for ocular research (Wisely et al., 2017). Chicks are diurnal and have a cone-rich retina with a high-density region, with parallels to the human macula. These similarities to the human eye make it attractive to model damaging retinal conditions and study the underlying cellular and molecular mechanisms relevant to neuroprotection (T. Heisler-Taylor et al., 2021).

## Materials and Methods

### Animals

This research adheres to the principles of the National Institutes of Health and the ARVO Statement for the Use of Animals in Ophthalmic and Vision Research. It was conducted under a protocol approved by The Ohio State University Institutional Animal Care and Use Committee. Newly hatched wild type leghorn chicks (Gallus gallus domesticus) were obtained from Meyer Hatchery (Polk, Ohio). Postnatal chicks were housed in a stainless-steel brooder at around 32℃ and kept on a 12-hour light, dark cycle. Water and Purina™ chick starter were provided ad libitum.

### Ocular Injections

Chicks were anesthetized via inhalation of 2-3% isoflurane in oxygen using a Kent Scientific VetFlo Calibrated Vaporizer. A 5% betadine antiseptic solution was administered on the ocular surface of each eye in addition to 10% betadine topical solution being applied to the superior eyelid and periocular region. The topical betadine was left to dry for at least 30 seconds before performing the injection through the superior palpebra into the globe with a 29 gauge insulin syringe. Once the injection was complete, the needle was held in place for a few seconds and then withdrawn. The eye was then rinsed with sterile saline. A 0.5% tetracaine ophthalmic drop solution was applied for topical anesthesia as needed.

### Ibudilast and AV1013 studies

Study drugs were obtained: ibudilast (InvivoChem, V3145), AV1013 (TRC, A618170), and NMDA (Sigma, M3262). The chicks were treated with intravitreal injections (20µl) of ibudilast (1x 10^-5^ mg/ml to 4.0 mg/ml) or AV1013 (1.0 mg/ml) in the left eye and control vehicle in the right eye under NMDA (25 mM) damage conditions. The experimental drug was also given 24 hours before damage for some experiment, including the ibudilast dose-testing, OCT, and ERG studies. Toxicity studies were conducted on undamaged chicks. Ibudilast was injected 2 hours after damage for the drug timing studies. Chicks were sacrificed one day after injection (D1) for cell death (TUNEL) and immunohistochemistry analysis, nine or ten days after injection (D9 or D10) for retinal layer thickness measurements and immunohistochemistry analysis, and seven to fourteen days after injection (D7 or D14) for SD-OCT and ERG studies.

### Enucleation, Fixation, and Sectioning

Chicks were euthanized and eyes enucleated. Eyes were bisected across the equator and the gel vitreous removed from the posterior eye cup. Eyes were fixed (4% paraformaldehyde plus 3% sucrose in 0.1 M phosphate buffer, pH 7.4, 30 min), washed two times in PBS, cryoprotected in PBS plus 30% sucrose, immersed in embedding medium (OCT-compound; Tissue-Tek), and freeze-mounted onto sectioning blocks. 12 µm transverse sections of the retina were cut and mounted onto SuperFrost Plus™ slides (Fisher Scientific). Sections from control and treated eyes from the same individual were placed consecutively on each slide to ensure equal exposures to reagents. Sections were air-dried and stored at − 20 °C until use.

### Terminal deoxynucleotidyl transferase dUTP nick end labeling (TUNEL)

TUNEL assay was performed to detect apoptosis and cell death with the In Situ Cell Death Kit (TMR red,

Roche Applied Science, #12156792910), as per the manufacturer’s instructions.

### Immunohistochemistry

Immunofluorescence was performed using primary antibodies (see antibody details, Table 1). Sections were permeabilized with 0.2% Triton X-100 and incubated with primary antibody solution (200 μl of antiserum diluted in PBS plus 5% normal goat serum, 0.2% Triton X-100, and 0.01% NaN3) overnight at 4 °C in a humidified chamber. The slides were then incubated for at least 1 h at room temperature in a humidified chamber with Alexafluor 488 conjugated secondary antibodies (Invitrogen). Omission of the primary antibody was used as a control for background staining. Cell nuclei were counterstained with DAPI (Invitrogen) or Hoechst 33342 (Cell Signaling Technology, #4082S). Slides were mounted in an 80% Glycerol solution.

**TABLE 1.** List of antibodies, working dilution, clone/catalog number, and source.

| **Antibody** | **Catalog number** | **Clone** | **Host** | **Dilution** | **Source** |
| --- | --- | --- | --- | --- | --- |
| CD45 | 8270-01 | Monoclonal/LT40 | Mouse | 1:500 | SouthernBiotech |
| pS6 | 5364 | Monoclonal/D68F8 | Rabbit | 1:750 | Cell Signaling Technology |
| pSIN1 | ABS1519 | Polyclonal | Rabbit | 1:200 | Millipore/Sigma |
| SIN1 | 15463-1-AP | Polyclonal | Rabbit | 1:100 | Proteintech |
| Islet-1 | 40.2D6 | Monoclonal/40.2D6 | Mouse | 1:100 | DSHB |
| PKCα | sc-80 | Monoclonal/MC5 | Mouse | 1:500 | Santa Cruz Biotechnology |
| Anti-rabbit IgG Alexa Fluor™ 488 | A-11008 | Polyclonal | Goat | 1:1000 | Invitrogen |
| Anti-mouse IgG Alexa Fluor™ 488 | A-11029 | Polyclonal | Goat | 1:1000 | Invitrogen |

### Single-cell RNA sequencing of retinas

In this study, we performed a secondary analysis of scRNA-seq libraries generated from NMDA-damaged chick retinas at 24hr, 48hr, and 72hr and saline controls which were originally generated and previously reported (Campbell, Blum, et al., 2021; Campbell, Fritsch-Kelleher, et al., 2021; Hoang et al., 2020; Palazzo, Deistler, Hoang, Blackshaw, & Fischer, 2020b). Two libraries were generated for each condition and evaluated for gene expression of MIF, putative MIF receptors, and PDE’s.

We also generated new libraries of undamaged and NMDA damaged chick retinas treated with ibudilast or AV1013 at 24hr. Retina samples were isolated from 12-14 week chicks. Four chick retinas were pooled for the library for each condition. Retinas were dissociated in a 0.25% papain solution in Hank’s balanced salt solution (HBSS), pH=7.4, for 30 min. To remove large particulate debris, dissociated cells were passed through a 70μm sterile filter. Cells were assessed for viability and diluted (700 cell/μl). Each single cell cDNA library was prepared, targeting 10,000 cells per sample. The cell suspension and Chromium Next GEM (Gel Beads-in-emulsion) Single Cell 3’ V3.1 dual index libraries (10X Genomics) were loaded onto chips to capture individual cells with individual gel beads in emulsion (GEMs) using 10X Chromium Controller. The optimal amplification for cDNA and library was 13 and 12 cycles, respectively. Novaseq6000 (Novogene) was used for sequencing with 150 paired end reads. Fasta sequence files were annotated, demultiplexed, and aligned using the chick ENSMBL database (GRCg7b, Ensembl release Gallus gallus 108) and 10x Genomics Cloud Analysis/ Cell Ranger software. Unique molecular identifier bar codes were utilized for gene expression, and gene-cell matrices were constructed. Using Seurat and Signac toolkits, uniform manifold approximation and projection for dimension reduction (UMAP) plots were generated from aggregates of multiple scRNA-seq libraries (Butler, Hoffman, Smibert, Papalexi, & Satija, 2018; Satija, Farrell, Gennert, Schier, & Regev, 2015). Seurat was used to construct scatter, violin, and dot plots. Wilcoxon rank-sum test with Bonferroni correction was used to assess significant differences for violin and scatter plots. SingleCellSignalR was used to assess potential ligand–receptor interactions between cells within scRNA-seq datasets (Cabello-Aguilar et al., 2020; Campbell et al., 2022). To perform gene ontology (GO) enrichment, analyzed lists of differentially expressed genes were uploaded to ShinyGO v0.80 (http://bioinformatics.sdstate.edu/go/).

Genes that were used to identify different types of retinal cells included the following, similar to previous reports (Campbell et al., 2023; El-Hodiri et al., 2023): (1) Resting Müller glia: *GLUL, RLBP1,GPR37L1*, (2) Activated Müller glia: *VIM, MDK, TGFB2,* (3) Müller glia progenitor cells: *PCNA, CDK1, TOP2A*, (4) microglia: *C1QA, C1QB, CCL4,* (5) ganglion cells: *THY1, RBPMS2, NEFM,* (6) amacrine cells: *CALB2, TFAP2A,* (7) horizontal cells: *PROX1, CALB2, NTRK1,* (8) bipolar cells: *VSX1, OTX2, GABRA1*, (9) ON bipolar cells: *TRPM1, ISL1,* (10) OFF bipolar cells: *GRIK1,* (11) cone photoreceptors: *CALB1, GNAT2, OPN1LW,* (12) rod photoreceptors: *RHO, NR2E3,* and (13) oligodendrocytes: *NKX2-2, OLIG2*.

### Data availability

Cell Ranger output files for Gene-Cell matrices for scRNA-seq data for libraries from the AV1013, ibudilast, NMDA+AV1013, and NMDA+ibudilast-treated retinas are available through GitHub: https://github.com/ccebulla/Chick_scRNAseq_libraries.git. The previously generated libraries from saline and NMDA-treated retinas which were previously published (Campbell, Blum, et al., 2021; Campbell, Fritsch-Kelleher, et al., 2021; Hoang et al., 2020; Palazzo et al., 2020b) are available through GitHub: https://github.com/jiewwwang/Single-cell-retinal-regeneration.git and https://github.com/fischerlab3140/scRNAseq_libraries.git. scRNA-seq datasets are deposited in GEO (GSE242796) and Gene-Cell matrices for scRNA-seq data for libraries from saline and NMDA-treated retinas are available through NCBI (GSM7770646, GSM7770647).

### Electroretinograms (ERG)

Electrophysiology was performed as previously described (T. Heisler-Taylor et al., 2021). Briefly, light adapted ERGs were recorded with the Celeris (Diagnosys, LLC) system on subsets of chicks (n=3/condition). Chicks were kept anesthetized as mentioned above via isoflurane. Genteal Severe gel was topically administered to the Celeris bright flash stimulators (5mm apparatuses) to moisten the cornea and facilitate electrical contact. Stimulators were aligned on each eye as per the manufacturer’s recommendations and both eyes were tested in a rapid alternating manner.

### Spectral domain optical coherence tomography (SD-OCT)

SD-OCT was performed on chicks as previously described (T. Heisler-Taylor et al., 2021). Briefly, an Envisu R2200 SDOIS with a rabbit bore lens was used to capture OCT b-scans of chick retina. Chicks were anesthetized as mentioned above via isoflurane. A small 2mm Barraquer wire eye speculum was used to hold the eyelid open, Genteal Severe gel was used to hydrate the eye not being actively imaged while Systane Ultra drops were used to hydrate the eye being actively imaged. The chick and probe were aligned so that the pecten could serve as a standard reference point in the constructed *en face* view.

### Image analysis

Photomicrographs were obtained using a Leica DM5000B fluorescent microscope and Leica DC500 digital camera or a Nikon Eclipse Ts2R fluorescent microscope with a panda sCMOS camera. Confocal images were obtained using a Nikon AXR confocal microscope. Treatment groups were analyzed in a masked fashion. To minimize the variability of region-specific differences within the retina, the same region of retina was evaluated for the fellow-eye control and treated areas. Identical illumination, microscope, and camera settings were used. The retina was selected as the region of interest from the 200X field of view. Image analysis was performed with ImageJ64 (ImageJ64, National Institutes of Health, Bethesda, MD, USA) as previously described or with NIS-Elements utilizing custom general analysis 3 (GA3) macros. Islet-1 staining was performed on D9 tissue and analyzed for the count of horizontal, bipolar, and cholinergic amacrine cells within the INL of NMDA damaged eyes with and without ibudilast and AV1013. Cell counts were normalized by the INL length. PKCα staining was performed on D9 tissue and compared between ibudilast-treated and untreated NMDA damaged eyes. The count of bright PKCα positive cells were measured per INL area. An angle deviation, θ, was calculated as the angle between the major axis of the green PKCα positive filaments and the perpendicular cross section of the INL medial axis. CD45 staining was performed on both D1 and D9 NMDA-damaged tissue with and without MIF inhibitor treatment. At D1, the mean CD45 positive area above a threshold (75 in an 8-bit image), the average cell count, the average form factor 1 (4π x Area / Perimeter^2^), and the average aspect ratio were assessed within the full retinal area. At D9, the mean CD45 positive area above a threshold and the average cell count were compared between conditions in the full retinal area, the outer retinal photoreceptors (PR-IS/OS), the mid retina (OPL-INL), and the inner retina (RNFL-IPL). pS6 and pSIN1 staining were both compared on D1 NMDA-damaged tissue with ibudilast and rapamycin treatments by assessing the mean intensity between paired subjects: NMDA±Rapamycin, NMDA±ibudilast, and NMDA+ibudilast±Rapamycin. TUNEL analysis was performed using ImageJ and NIS-Elements. Multichannel thresholding was performed to quantitate TUNEL-positive cells that also exhibited DAPI positivity in ImageJ (Tyler Heisler-Taylor et al., 2018) and a similar macro was designed in NIS-Elements. Conditions were compared as either the TUNEL positive cell density or as a ratio between paired eyes.

### Statistics

Data were collected on standardized tables after analysis in a masked fashion. Error bars are listed with standard deviation. Paired student’s T-tests, Wilcoxon paired non-parametric tests, and 1-way ANOVA tests with Tukey post-hoc corrections were performed as appropriate in JMP and GraphPad PRISM 10 to evaluate the differences between experimental groups, with p≤0.05 considered statistically significant. scRNA-seq data analysis used Wilcoxon Rank Sum test with Bonferroni correction.

## Results

### scRNA-seq analysis of MIF and PDE pathways show active modulation during excitotoxic retinal damage

We began by assessing patterns of expression of players in the MIF and PDE pathways across different cell types in NMDA damaged and control retinas using single cell RNA sequencing (scRNA-seq) analysis. We performed secondary analysis of scRNA-seq libraries of the chick retina which were originally generated and described in previous studies (Campbell, Blum, et al., 2021; Campbell, Fritsch-Kelleher, et al., 2021; Hoang et al., 2020; Palazzo et al., 2020b).

scRNA-seq libraries of retinas treated with NMDA at 24, 48, and 72 hrs were analyzed and compared to saline controls. Eight independent sets of libraries were generated, aggregated, and re-analyzed. After processing to remove dying cells, doublets, and cells with low gene/cell ratios for a total of more than 90,000 cells. Uniform Manifold Approximation and Projection (UMAP) plots were generated, and discrete clusters of different retinal cell types were identified based on gene expression of well-established markers, as described in methods. We created dot plots, UMAP feature plots, and violin plots to examine the libraries for expression changes and patterns in *MIF*, its receptors *CD44* and *CD74*, and several isoforms of PDE that are targeted by ibudilast (i.e., *PDE3A, 4B, 4D, 10A, 11A*) (Fig 1C-L, Table S1). *MIF* is expressed to some degree in all retinal cells, with highest levels of expression in MG, ganglion, amacrine, bipolar, and horizontal cells (Fig 1C). *MIF* expression goes up in all retinal cell types post damage, especially 24 hr post NMDA (Fig 1K, L, S1). MIF receptors *CD74* and *CD44* were highly expressed in microglia, with modest *CD44* expression also in activated MG post damage (Fig 1D, E).

**Fig 1:**
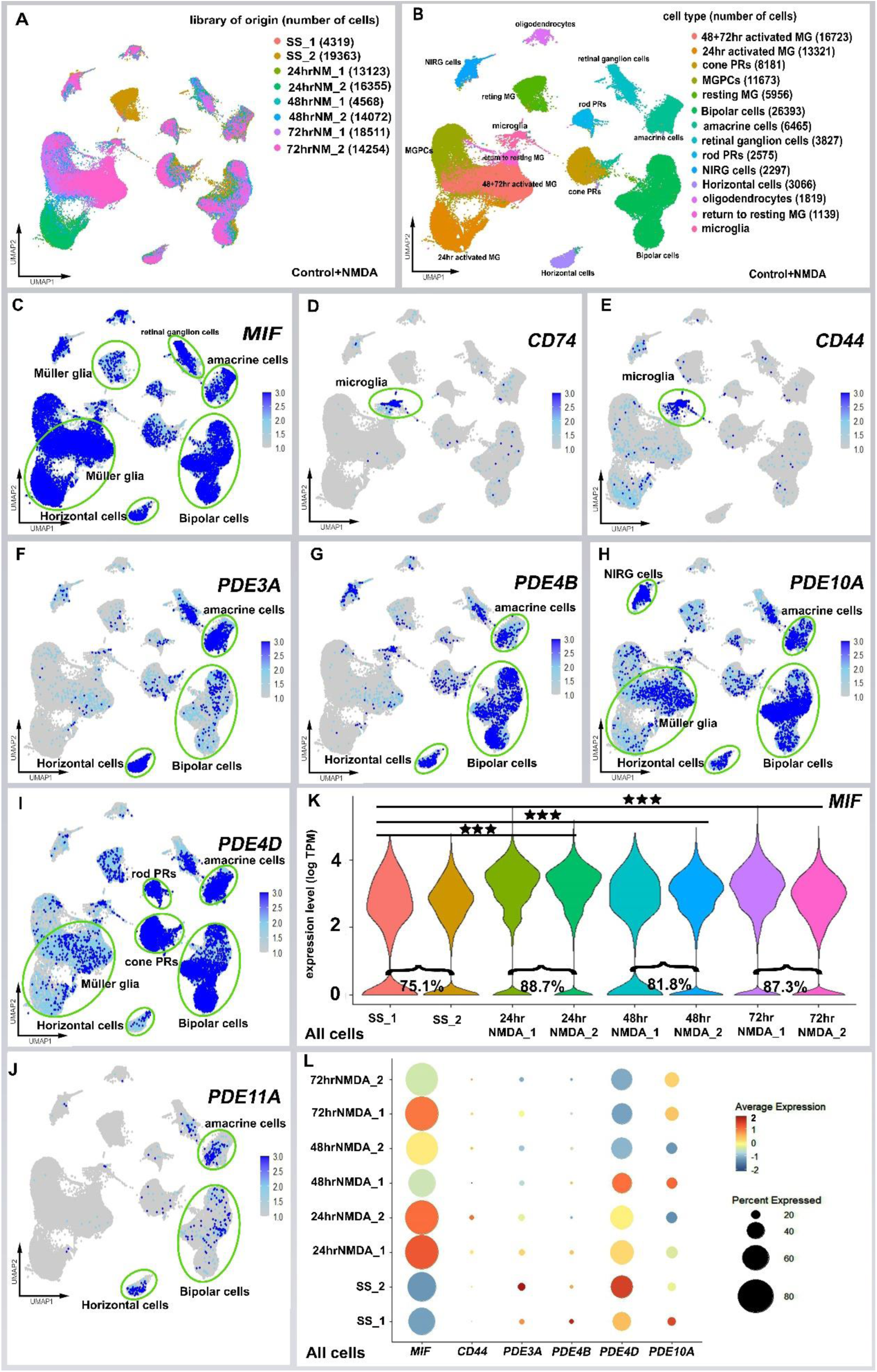
Upregulation of MIF and PDE-related gene expression in damaged chick retina. UMAPs of aggregated scRNA-seq libraries from two independent isolations prepared from chick control (Saline) and NMDA-damaged retinas at 24hr, 48, and 72hr (A). Clusters of different types of retinal cells were identified based on collective expression of different cell-distinguishing markers (B). UMAP plots illustrate expression of *MIF, CD74, CD44, PDE3A, PDE4B, PDE4D, PDE10A, PDE11A* (C-J). Violin plots illustrate percentage and expression levels of MIF in control and NMDA-damaged retinas within all retinal cells (K). Dot plot shows distinct patterns of gene expression of MIF and PDE genes in all retinal cells of control and NMDA damaged chick retinas (L). Significance of difference was determined by using a Wilcoxon rank sum with Bonferroni correction (***p < .001).

Many PDE isoforms were differentially expressed in retinal neuronal cell populations and actively modulated during retinal damage. All PDE’s examined were expressed in populations of bipolar cells, with particularly high levels of expression of *PDE4B, PDE4D*, and *PDE10A* (Fig 1F-J, L). *PDE3A* expression was notable for very strong expression in horizontal cells, as well as in amacrine cells and a subset of retinal ganglion cells (RGC) (Fig 1F); *PDE10A* had a similar pattern of expression in these neurons (Fig 1H). *PDE4D* had high expression in amacrine cells, and some RGCs, but lower expression in horizontal cells than *PDE3A* (Fig 1I). *PDE4D* was the only PDE in this group that was highly expressed in the rods and cones. The MG predominantly expressed *PDE4D* and *PDE10A*, which was highest in the 48-72h activated MG population (Fig 1H, I). Microglia expressed predominantly *PDE4B, PDE4D*, and *PDE10A*. *PDE11A* had the lowest expression levels in the retina, mostly in the horizontal cells as well as RGC, amacrine, and bipolar cells (Fig 1J).

To gain further insight on the *MIF* and *PDE* gene expression of the MG and microglia populations, we isolated these cell subsets and re-embedded them in UMAP plots, created violin plots, and ran statistical analyses of the pooled libraries (Fig 2). *MIF* was more highly expressed in MG than in microglia (Fig 2 C, H), while MG had no *CD74* and only some *CD44* after damage (Fig 2D, E, I, J). The MG showed significant increases in expression of *MIF*, particularly 24 hours after NMDA treatment, compared to levels seen in resting and undamaged MG (Fig 2C, H). Similarly, microglial *MIF* levels were upregulated, particularly 24h after damage. *PDE4D* and *PDE10A* were predominantly expressed in the MG, rather than microglia, and expression of both increased significantly 24h post NMDA damage. *PDE10A* further increased at 48-72h in the MG (Fig 2F, G, K, L). The data from these studies indicate that there are multiple populations of cells that could respond to *MIF* and *PDE* inhibition with ibudilast treatment and provide a rationale to further investigate ibudilast’s ability to ameliorate retinal excitotoxic damage.

**Fig 2:**
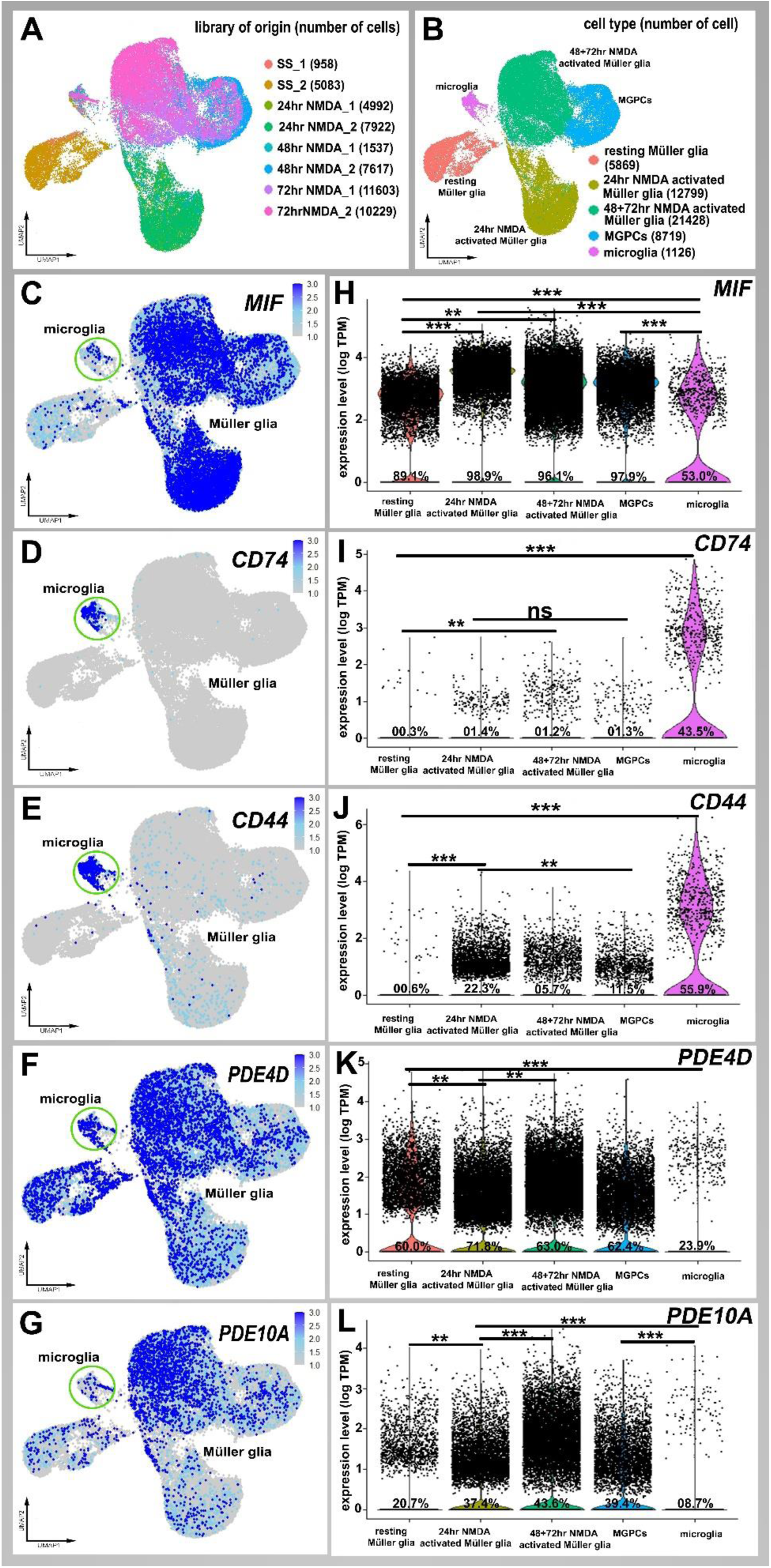
scRNA-seq analysis of MIF and PDE-related genes MG and microglia. UMAPs of aggregated scRNA-seq libraries prepared from 2 independent chick libraries of control (saline) and NMDA-damaged retinas at 24hr, 48, and 72hr (A). Clusters of MG and microglia were identified based on collective expression of different cell-distinguishing markers (B). scRNA-seq was used to identify patterns and levels of expression of MIF and its receptor in chick control (saline) and NMDA-damaged retinas at 24hr, 48, and 72hr. UMAP heatmap plots (feature plots (C-G)) and violin plots (H-L) illustrate patterns and levels of expression for *MIF, CD74, CD44, PDE4D, PDE10A* genes in MG and microglia. \*\**P*<1×10^−5^, \*\*\**P*<1×10^−10^ (Wilcoxon Rank Sum Test with Bonferroni correction)

### Ibudilast and its analog AV1013 do not induce toxicity in the chick retina

We investigated the potential retinal toxicity of ibudilast and its analog AV1013 in undamaged chick eyes. Intravitreal delivery of the maximum soluble ibudilast in sterile water (4mg/mL) did not show toxicity on ERG (Fig S1A-E), TUNEL assay (Fig S1F), and retinal layer thickness measured by spectral domain optical coherence tomography (SD-OCT, (Fig S1G)). Similarly, AV1013 was evaluated on TUNEL analysis in undamaged eyes and resulted in no cell death (Fig S1F). NMDA damaged eyes were compared as a positive control (Fig S1F). In undamaged eyes treated with ibudilast or AV1013, intraocular pressures were not significantly different from controls (Fig S2A-C). The drug injections did not alter growth rates of the animals (Fig S2D).

### Ibudilast protects against neuronal cell death during excitotoxic damage, while analog AV1013 does not

The increase in MIF and PDE expression in neurons and MG after damage suggests that these proteins may be involved in cell responses to injury and that an inhibitor that targets both pathways can modulate the cellular response to damage. Given the lack of toxicity with intravitreal injection, we investigated the ability of ibudilast and its analogue AV1013 to prevent NMDA-induced injury in the retina. To that end, chick eyes were injected with ibudilast (20uL of 4mg/mL) along with NMDA (Fig 3) and numbers of dying cell were quantified using TUNEL analysis (Fig 3A-D). Notably, the chick retinal ganglion cells are resistant to NMDA-induced excitotoxicity, unlike retinal ganglion cells in the mammalian retina; the dying cells localize to the INL in the chick (Fischer & Reh, 2003; Sattayasai, Zappia, & Ehrlich, 1989). Ibudilast significantly reduced dying cells in the inner nuclear layer (INL).

**Fig 3.**
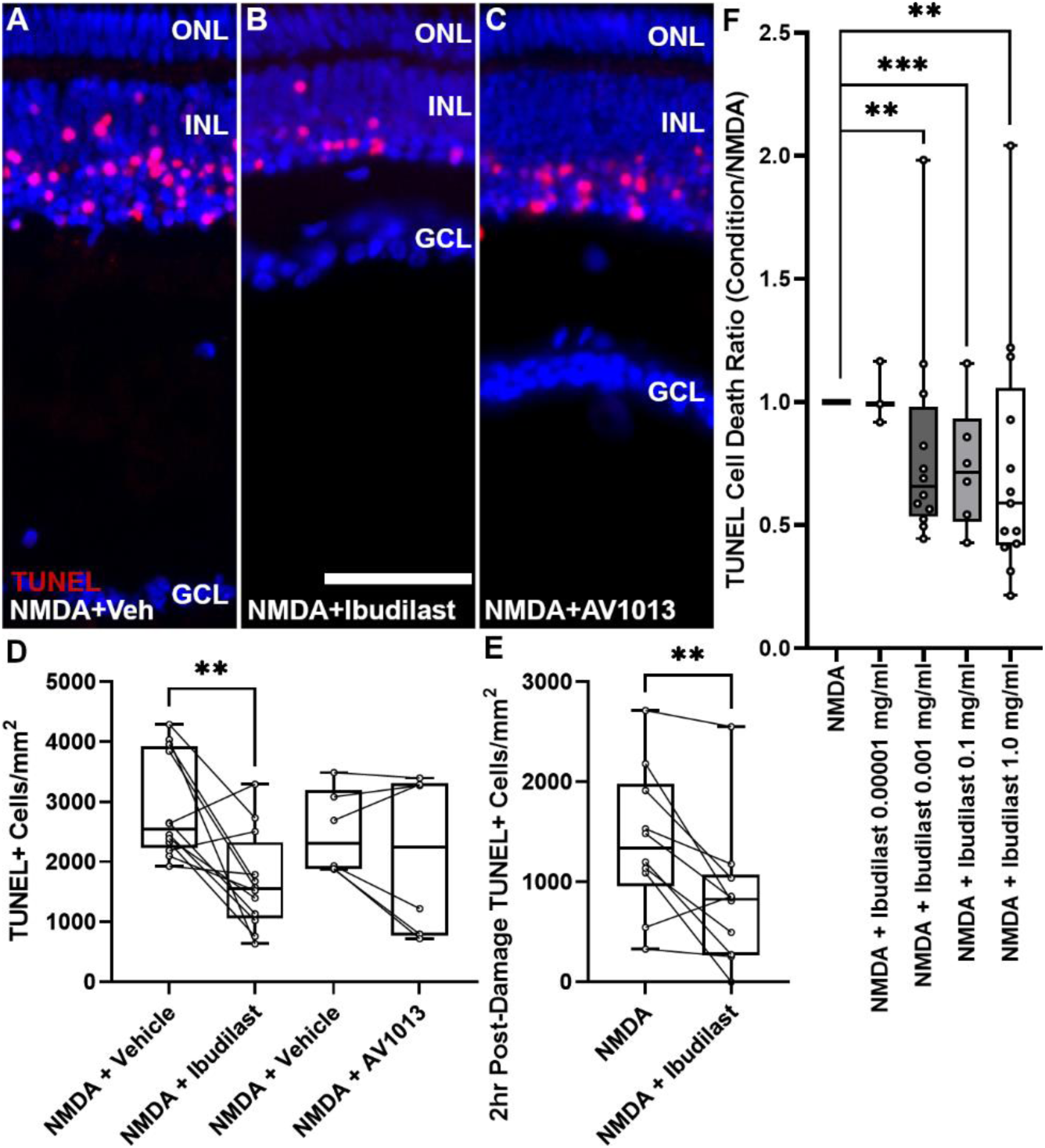
Ibudilast protects against neuronal cell death during excitotoxic damage, while analog AV1013 does not. Representative TUNEL images of retinas from eyes 24h post injection with NMDA only (A) NMDA+ibudilast (B) or NMDA+AV1013 (C) to quantify dying cells. The graph in (D) represents the number of TUNEL-positive cells (±SD) for each treatment group. Ibudilast also significantly reduced the number of TUNEL-positive cells when injected 2 hours *after* NMDA damage induction (E). Ibudilast’s dose-dependent effect is illustrated in graph (F). Each dot represents one biological replicate. Significance of difference (**p<0.01, ***p<0.001) was determined by using a paired t-test comparing the fellow control eye and the Steel-Dwass test with a control for the dose-dependent effect. The calibration bar (50 μm) in panel B applies to A-C. Abbreviations: ONL – outer nuclear layer, INL – inner nuclear layer, GCL – ganglion cell layer.

Further, we observed a dose-dependent decrease in TUNEL positive cells in the INL with ibudilast treatment (Fig 3F). The neuroprotective effects of ibudilast started at a concentration as low as 1 ug/mL and reached a maximum effect at 1 mg/mL (Fig 3F). The maximum effect for ibudilast exceeded 40% reduction in TUNEL positive cells, compared to NMDA control (Fig 3D). Interestingly, AV1013, the MIF inhibitor that lacks the PDE inhibitor activity, did not prevent NMDA-induced neuronal cell death (Fig 3B,D), which suggests the possible importance of PDE inhibition for ibudilast’s neuroprotective effect.

To further characterize the potential clinical utility of ibudilast we investigated whether ibudilast could impact dying neurons *after* damage had been initiated. We injected ibudilast 2 hours post NMDA damage. Ibudilast treatment significantly reduced TUNEL positive dying cells 41% compared to vehicle (Fig 3E), indicating a proof of concept that the drug could help salvage neurons after damage had occurred.

### Ibudilast protects against excitotoxic damage-induced thinning of retinal layers

We next assessed whether ibudilast or AV1013 could preserve retinal structure by performing SD-OCT analysis to quantitate the thickness of the retinal layers. The death of neurons from chick NMDA damage in the inner retina leads to retinal thinning, which we previously demonstrated could be measured with SD-OCT (T. Heisler-Taylor et al., 2021). We evaluated retinal layer thickness 10 days post NMDA treatment, which showed that ibudilast, but not AV1013, significantly preserved retinal thickness (Fig 4A-D, NMDA 246.5±5.044 microns vs. NMDA+ibudilast 259.6±3.215, p=0.0260). Moreover, we previously identified a hyperreflective band in the inner nuclear layer occurring 10 days after NMDA damage (T. Heisler-Taylor et al., 2021). Areas of inner retinal hyperreflectivity on OCT have been noted clinically in humans with acute retinal ischemia and are thought to represent edema of cells and nerve fibers; activated microglia are also hypothesized to compose this band (Feucht et al., 2018; Wenzel et al., 2022). Interestingly, this hyperreflectivity was less prevalent in the ibudilast-treated eyes (Fig 4A-C).

**Fig 4:**
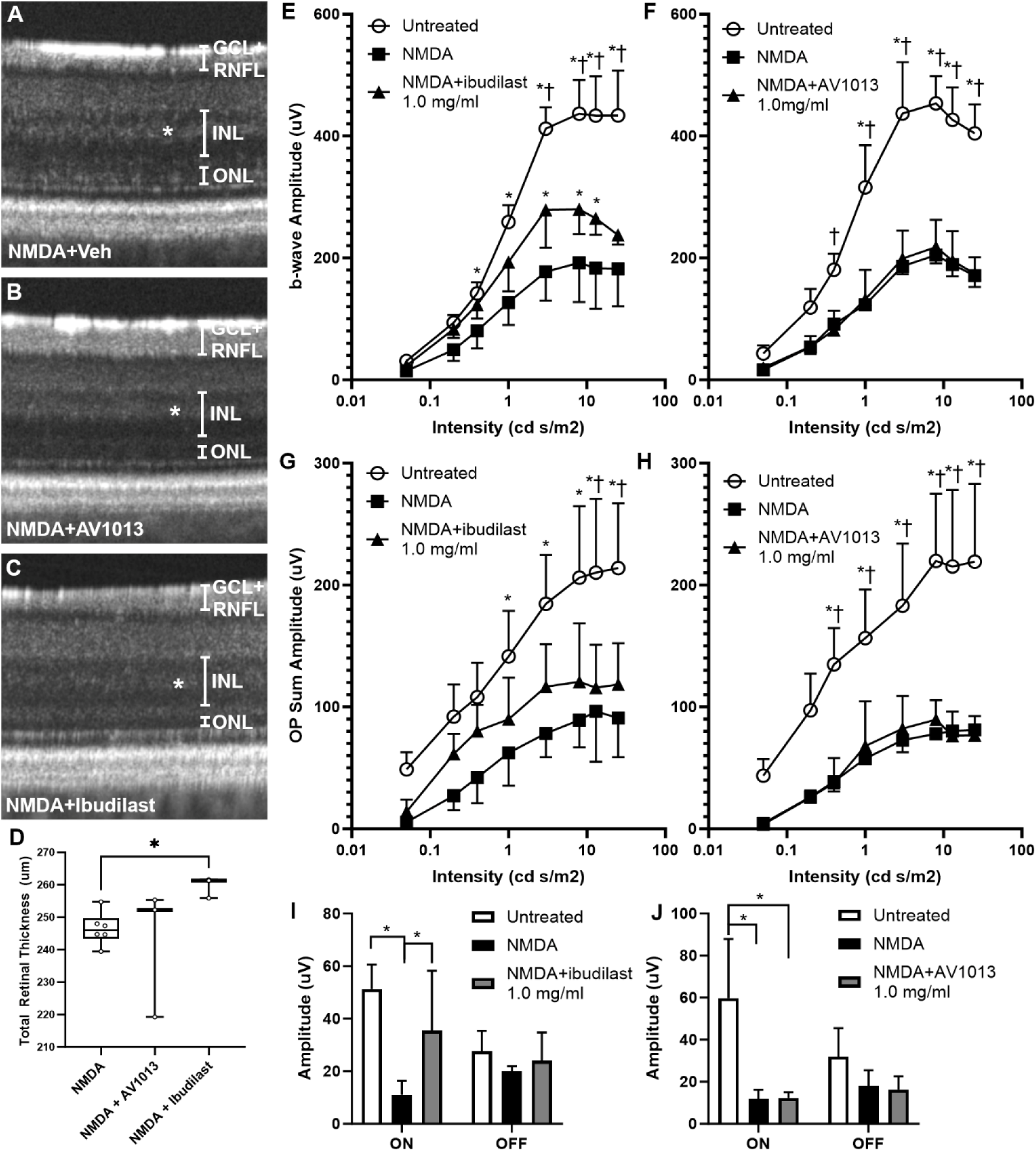
MIF inhibition with ibudilast preserves NMDA-damaged chick retinal thickness on SD-OCT and bipolar cell function by ERG. Representative SD-OCT b-scans of chick eyes D10 post injection with NMDA (A), NMDA+AV1013 (B), or NMDA+ibudilast (C). The * indicates the hyper-reflective band in the INL. The graph in (D) represents total retinal thickness of the 3 groups above (±SD) (n=3 chicks/group, p=0.0260, Dunn’s multiple comparison test). Light adapted ERG results (n=3/group) are shown in panels (E-J). b-waves amplitudes show significant preservation with ibudilast during NMDA damage (E), while AV1013 does not show the same protection (F). Oscillatory potentials show a similar pattern of improvement with ibudilast treatment (G) but not with AV1013 treatment (H), although ibudilast’s effect was not significant from NMDA alone. The long-flash ERG ON responses were significantly preserved with ibudilast (I) while AV1013 did not show any functional recovery (J). * indicates p<0.05 compared with NMDA damaged eyes, † indicates p<0.05 compared with NMDA+Drug injection eyes. Abbreviations: RNFL—retinal nerve fiber layer, IPL—inner plexiform layer, INL—inner nuclear layer, ONL—outer nuclear layer, GCL-ganglion cell layer

### Ibudilast preserves retinal function in excitotoxic damage

In addition to studying preservation of retinal anatomy, we investigated ibudilast and AV1013’s ability to preserve retinal function following NMDA damage. We performed light adapted electroretinogram (ERG) analysis after NMDA damage to evaluate retinal function and to provide insights on the specific cell populations that may be benefitted by drug treatment. Specifically, the ERG b-wave, which represents bipolar and MG cell function, was significantly preserved by ibudilast, but not by AV1013 (Fig 4E, F). Amacrine cells are the major cell type impacted by NMDA damage in the chick retina (Fischer, Seltner, Poon, & Stell, 1998). Amacrine cells synapse with bipolar cells in the inner plexiform layer (IPL) and influence their function. Oscillatory potentials are used to assess the function of inner retinal cells, including amacrine cells on ERG. NMDA treatment led to large reduction in oscillatory potential amplitudes that was not significantly recovered by ibudilast treatment (Fig 4G, H).

Using the ERG long flash ON/OFF response test, which allows for specific characterization of the functionality of the ON (depolarizing) and OFF (hyperpolarizing) bipolar cells, we further found that the ON bipolar cells are uniquely susceptible to NMDA damage compared to OFF bipolar cells, consistent with reports in the literature (T. Heisler-Taylor et al., 2021). Ibudilast treatment significantly rescued the function of ON bipolar cells and restored ON-wave amplitudes (Fig 4I, J). In contrast, ibudilast did not preserve OFF bipolar cell function (Fig 4I, J).

### Bipolar cell preservation by Ibudilast

Since our ERG results showed that during NMDA damage ibudilast specifically preserved the ON bipolar cell pathway function, we sought to validate our findings with IHC using bipolar cell specific markers.

Islet-1 is expressed by ON bipolar cells, and is also expressed by about half of the horizontal, cholinergic amacrine cells, and ganglion cells in the chick retina (Fischer, Foster, Scott, & Sherwood, 2008; Ritchey et al., 2010). These four cell populations can be differentiated by their localization; horizontal cells are a linear array of cells that are close to the outer retina, bipolar cells are localized to the middle of the INL, cholinergic amacrine cells at the proximal border of the INL, and ganglion cells in the GCL (Fig 5D). Immunolabeling for Islet-1 showed that ibudilast, but not AV1013, significantly preserved the ON bipolar cells damaged by NMDA treatment (Fig 5A-E). Interestingly, the Islet-1 expressing horizontal cells also showed significant preservation in the Ibudilast treated group compared to NMDA alone (Fig 5E).

**Fig 5.**
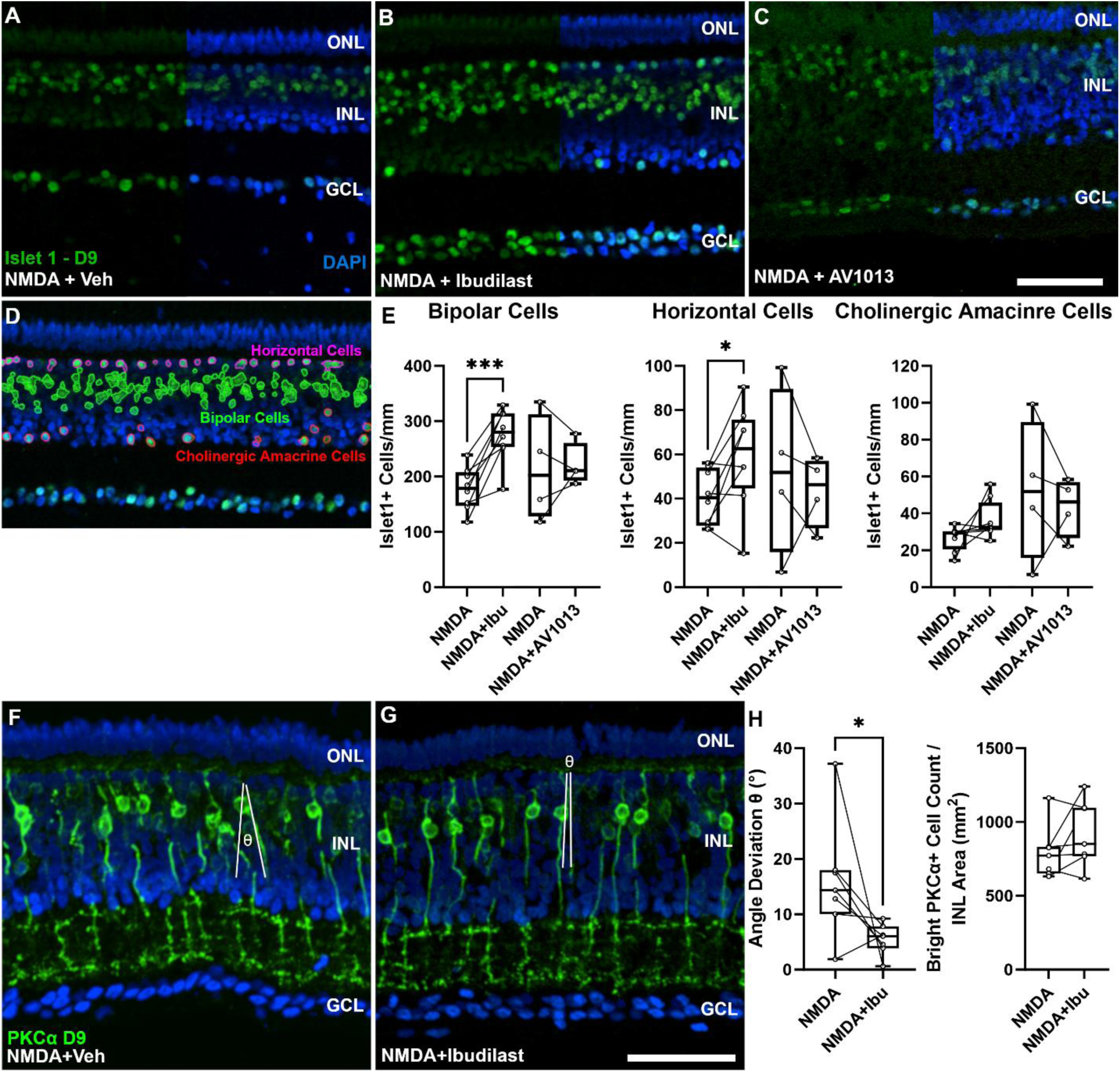
Ibudilast preserves NMDA-damaged chick retinal ON bipolar and horizontal cells. Representative images of retinas from eyes injected with NMDA only (A), NMDA+ibudilast (B), or NMDA+AV1013 (C) at D9 and stained for islet1 (green) with DAPI nuclear counterstain (blue, merged only in the right half of the image). Islet1 stains ON bipolar cells but can also detect horizontal and cholinergic amacrine cells using anatomic localization within retinal layers (NIS elements software segmentation (panel D)). The graphs in (E) represent the number of islet1 positive cells (±SD) for each cell type and treatment group. Ibudilast significantly preserved bipolar cells (*** represents p=0.0005) and horizontal cells (* represents p=0.0357) but not cholinergic amacrine cells (p=0.116) (F), Paired T-test is used for significance of difference calculations. AV1013 does not show any significant changes from NMDA. PKCα is used to stain rod bipolar cell (green) with DAPI nuclear counterstain (panels F-H). NIS software analysis is applied to quantify the integrity of the cells using angle deviation measures of PKCα positive cells and PKCα cell counts (H). Scale bar in D denotes 50 μm and applies to all images.

We further evaluated PKCα, which is expressed by rod bipolar cells and cone ON bipolar cells (Ritchey et al., 2010). PKCα positive cell numbers and staining area did not change with ibudilast treatment during NMDA damage (Fig 5F-H). However, the rod bipolar cell integrity appeared to be qualitatively preserved and there was less deviation of the axonal angles with ibudilast treatment (Fig 5H).

### Ibudilast and AV1013 largely do not influence the accumulation of microglia in the retina after damage

Excitotoxic damage triggers microglia activation and recruitment to the retina (Palazzo, Deistler, Hoang, Blackshaw, & Fischer, 2020a; Palazzo et al., 2022; Silverman & Wong, 2018; Wang & Wong, 2014). We sought to evaluate the effects of ibudilast on microglia distribution and morphology in the retina after NMDA damage. Microglia were found in the IPL 24h after NMDA damage (Fig 6A). The distribution was not altered by treatment with either ibudilast or AV1013 at this timepoint (Fig 6B, C). There was no significant difference in CD45 intensity or area above a threshold (Fig 6D). When measured 9 days after damage CD45 positive cells appeared throughout the retinal layers, including in the photoreceptor layer (Fig 6E). There was a significant decrease in the area and mean count of CD45+ cells in the photoreceptor layer with AV1013 treatment (Fig 6G, H) but no significant difference appeared in CD45+ cells in that layer with ibudilast treatment (Fig 6F,H).

**Fig 6.**
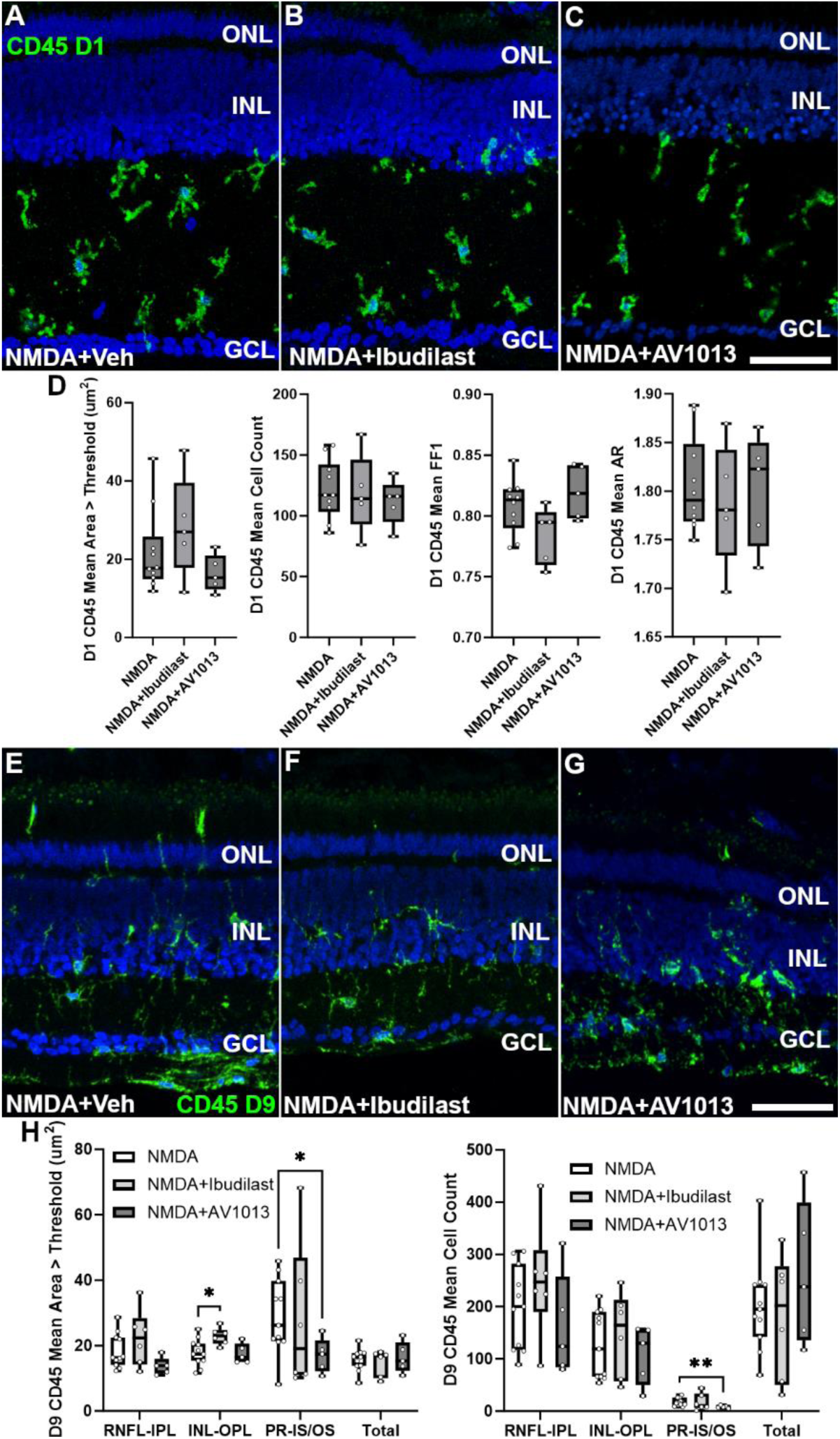
Ibudilast and AV1013 do not inhibit the accumulation of microglia in the retina after damage. Representative images of CD45 stained retinal sections from eyes post injection with NMDA (A, E), NMDA+ibudilast (B, F), or NMDA+AV1013 (C, G) at D1 (A, B, C) or D9 (E, F, G). Quantification of CD45 intensity, number of CD45+ microglia in the retina, and microglia activation (measured by analyzing morphological changes through FF1 and AR) are shown for the different treatment groups ((D), D1; (H), D9). The microglia are confined primarily to the inner plexiform layer (IPL) on D1 and there is no significant difference in any of the measures between the different treatments. At D9 (E-H) microglia are detected in all retinal layers. There is a slight statistically significant decrease in the area and mean count of CD45+ cells in the photoreceptor layer with AV1013 treatment ((H), * p<0.05, **, p<0.01) and a trend towards fewer CD45+ cells in the photoreceptor layer with ibudilast treatment, but this did not reach statistical significance (p=0.0659).

The morphology of microglia reflects the phenotype and activation status, with activated microglia displaying a more amoeboid shape and quiescent microglia a more ramified shape (Silverman & Wong, 2018). We performed aspect ratio (AR) and form factor 1 (FF1) analyses to quantitate potential morphology changes in microglia with ibudilast and AV1013 treatment (Fig 6D, H and S3). There was no difference in microglia morphology induced by either drug (Fig 6D, H).

### Ibudilast upregulates mTORC1 and mTORC2 gene expression in the MG and bipolar cells during damage

To further elucidate the mechanism of ibudilast’s protection of retinal neurons we conducted scRNA-seq analyses to probe genome-wide, cell-level transcriptomic changes. We created scRNA-seq libraries of retinas 24h post treatment with ibudilast or AV1013, with or without NMDA (Fig 7A-B). Libraries were generated, aggregated, and processed as described in methods. Distinct populations of retinal cells were identified based on expression of cell-distinguishing markers. Undamaged MG treated with ibudilast or AV1013 composed a distinct UMAP cluster of resting MG, while NMDA-treated retinas combined with those drugs produced a distinct cluster of activated MG (Fig 7A, B). Large numbers (>80,000 cells) of neurons and MG were isolated, while only a small number (559 cells) of microglia were captured. To evaluate important signaling pathways during damage impacted by ibudilast, compared to AV1013, gene ontology (GO) enrichment analysis of differentially expressed genes was performed. In the damaged MG, ibudilast significantly upregulated gene modules associated with many signaling pathways including Hippo, PI3K-Akt, mTOR, and HIF-1 (Fig 7C).

**Fig 7:**
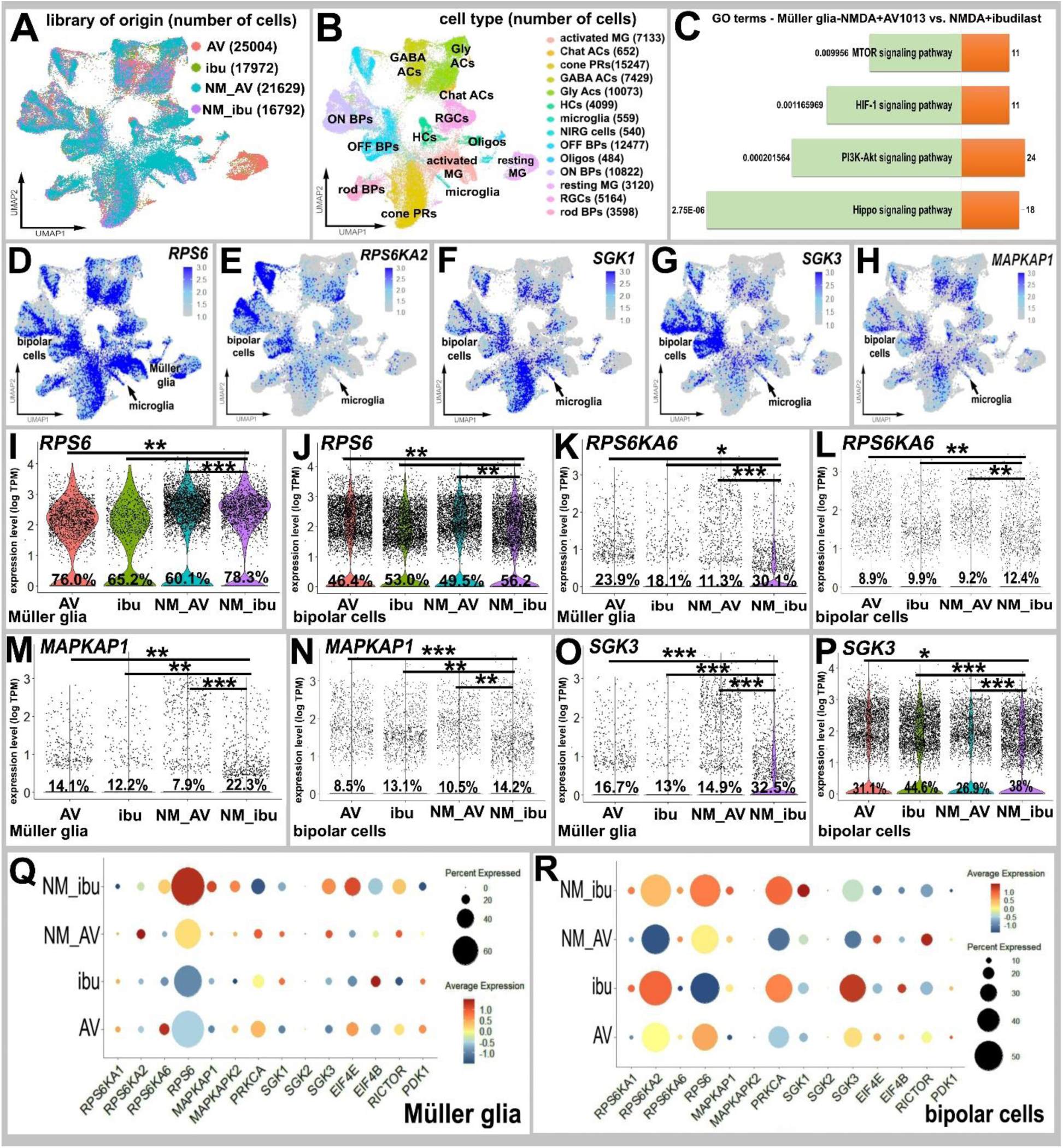
Upregulation of mTOR pathway genes in chick retina treated with NMDA+ibudilast compared to NMDA+AV1013. UMAPs of aggregated scRNA-seq libraries prepared from undamaged or NMDA damaged chick retinas treated with AV1013 or ibudilast (A). Clusters of retinal cells were identified based on collective expression of different cell-distinguishing markers (B). ShinyGO KEGG Pathway MG enrichment analysis was performed for upregulated pathways (green) and associated p-value, and the number of enriched genes in each pathway category (orange) are displayed (C). UMAP feature plots illustrate expression of *RPS6, RPS6KA2, SGK1, SGK3* and *MAPKAP1* (D-H). Violin plots illustrate expression levels of *RPS6* (I, J), *RPS6KA6* (K, L), *MAPKAP1* (M, N), *SGK3* (O, P) in MG and bipolar cells. Dot plots indicate the percentage of expression (size) and expression level (heatmap) for genes related to the mTORpathway (Q, R). (*P<0.05, **P<1×10−5, ***P<1×10−10, Wilcoxon Rank Sum Test with Bonferroni correction).

mTOR is a clinically relevant, druggable pathway, with long established therapeutics available to target it (Wong, 2013; Yang et al., 2021). Thus, we probed the ibudilast/AV1013 scRNA-seq libraries for several players in the mTOR pathway to further investigate ibudilast’s impact on mTOR signaling. Notably UMAP feature plots showed expression in many retinal cell types of both mTORC1 and mTORC2 pathway genes, including *RPS6*, *RPS6KA2* (mTORC1), *SGK1/3, MAPKAP1,* and *PRKCA* (mTORC2) (Fig 7D-R, Fig S4, Table S2, S3). We further focused on analyzing the impact on gene expression in cells likely mediating ibudilast’s survival effect, i.e., bipolar cells and MG. Interestingly, both the MG and bipolar cells showed high levels of expression of many of these mTOR pathway genes. Ibudilast induced a significant increase in expression of *RPS6* in both MG and bipolar cells compared to AV1013 during damage (Fig 7I, J). Similarly, mTORC1 pathway gene *RPS6KA6* was significantly upregulated in damaged MG and bipolar cells by ibudilast treatment (Fig 7K, L, Q, R) while mTORC2 genes *MAPKAP1* (Fig 7M, N, Q, R), *SGK3* (Fig 7O, P, Q, R), *MAPKAP2*, and *EIF4E* showed moderate upregulation, mostly in the MG (Fig 7Q). *SGK1* and *PRKCA* were also upregulated with ibudilast treatment in the MG and particularly in the bipolar cells (Fig S4). A schematic of the genes upregulated with ibudilast treatment during retinal damage are shown (Fig 10).

### Validation of ibudilast’s impact on the mTOR signaling pathway

Our scRNA-seq analysis identified multiple mTOR pathway genes that are upregulated with ibudilast during damage. Some mTORC1 components had particularly strong upregulation, most notably *RPS6*, which encodes S6, a downstream target of mTORC1 (Laplante & Sabatini, 2012a). The upregulation of *RPS6* was observed in both MG and bipolar cells (Fig 7I, J). Our study also identified *MAPKAP1* upregulation by ibudilast. *MAPKAP1* encodes the protein, SIN1, which is a component of the mTORC2 complex. To investigate whether ibudilast increases mTORC1 signaling activity during damage, we performed confocal analysis of levels of phosphorylated S6 (pS6) in retinas treated with ibudilast under damage conditions compared to NMDA controls. In the chick retina, pS6 is known to be transiently upregulated in MG following NMDA-treatment, and this upregulation is blocked by rapamycin (Zelinka et al., 2016). Interestingly, ibudilast significantly increased pS6 levels throughout the retina, especially in the INL, compared to NMDA (Fig 8A, B, E). Rapamycin, an inhibitor of the mTORC1 pathway, blocked this increase (Fig 8C, D, F). We also evaluated SIN1 levels and phosphorylation status with ibudilast treatment during damage. While total SIN1 protein levels did not change (not shown), pSIN1 levels significantly increased with ibudilast treatment during damage (Fig 8G, H, K). Rapamycin also blocked ibudilast’s upregulation of pSIN1 in our damage model (Fig 8I, J, L).

**Fig 8.**
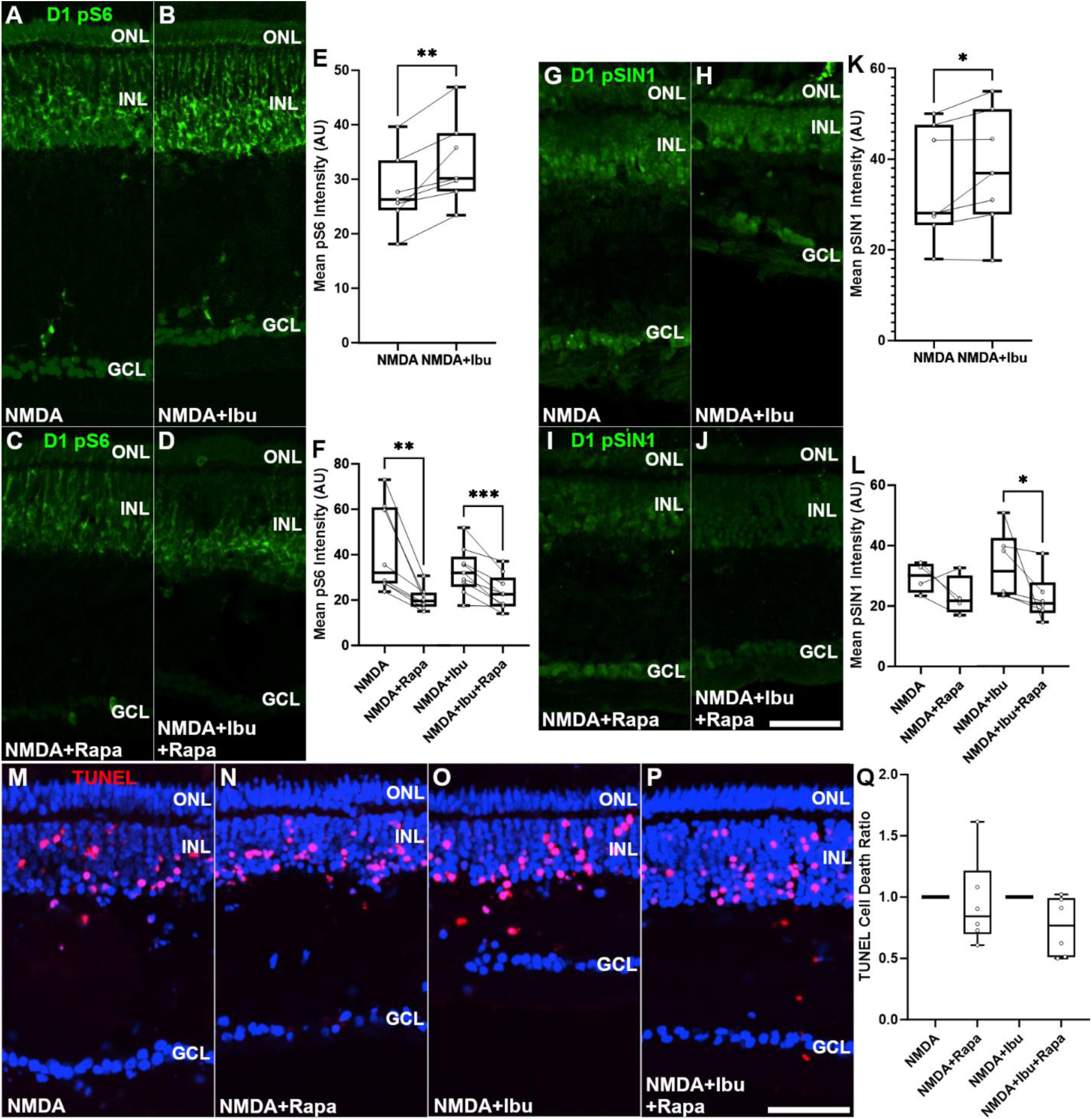
Validation of ibudilast’s impact on the mTOR signaling pathway and mTOR inhibitor rapamycin. Representative images of retinal sections from eyes injected with NMDA (A, G, M), NMDA+Rapamycin (B, H, N), NMDA+ibudilast (C, I, O), or NMDA+ibudilast+Rapamycin (D, J, P) at D1. Sections were stained for pS6 (A, B, C, D), pSIN1 (G, H, I, J), or cell death via TUNEL assay (M, N, O, P). Quantification of mean intensity between paired groups are shown for both proteins. Ibudilast treatment of NMDA damage was shown to elevate staining for both pS6 (E) and pSIN1 (K). Rapamycin treatment of NMDA damage was shown to inhibit pS6 staining both with and without ibudilast treatment (F). Rapamycin treatment of NMDA damage was shown to inhibit pSIN1 staining with ibudilast treatment (L). No differences were found due to rapamycin treatment of NMDA damage with or without ibudilast treatment (Q). (*p<0.05, **p<0.01, ***p<0.001)

### mTOR activation is not essential for ibudilast’s neuroprotective effect

We hypothesized that mTOR signaling might play a role in ibudilast’s neuroprotection of bipolar cells. To test that hypothesis, we used rapamycin to block the mTOR pathway and evaluated the ability of ibudilast to protect neuronal cells under damage conditions. As expected, rapamycin significantly decreased pS6 levels (Fig 8C,D) but it did not alter ibudilast’s protective effect during damage (Fig 8M-Q). This is in line with our previous results in NMDA damaged chick retina, where rapamycin had no effect on retinal cell death (Zelinka et al., 2016).

### Ligand receptor interaction analysis

To gain further understanding of the mechanism of ibudilast’s protection of ON bipolar cells we investigated putative autocrine and paracrine ligand receptor interactions between the MG and bipolar cell populations. We bioinformatically isolated MG and bipolar cells, re-embedded them in UMAP plots (Fig 9A-C), and probed for cell signaling networks and putative ligand-receptor (LR) interactions using SingleCellSignalR (Cabello-Aguilar et al., 2020). We compared ibudilast treated cells to AV1013 treated cells during NMDA damage to identify pathways changed by ibudilast treatment. The largest number of LR interactions was seen in the MG, acting in an autocrine fashion (Fig 9J-K). Further analysis extended to MG ligands acting on bipolar cell receptors in paracrine fashion from each treatment group and the most significant LR interactions were identified (Fig 9L, M). The top 40 LR interactions were the same between NMDA-ibudilast and NMDA-AV1013 treatment groups, except for LRPAP1-LRP8/SORL1 and TGFB2-TGFBR1, which were unique to the NMDA-ibudilast treatment group, and GNB3-GABBR2, NXPH1-NRXN3, and PTN-PTPRZ1, which were unique to the NMDA-AV1013 treatment group.

**Fig 9.**
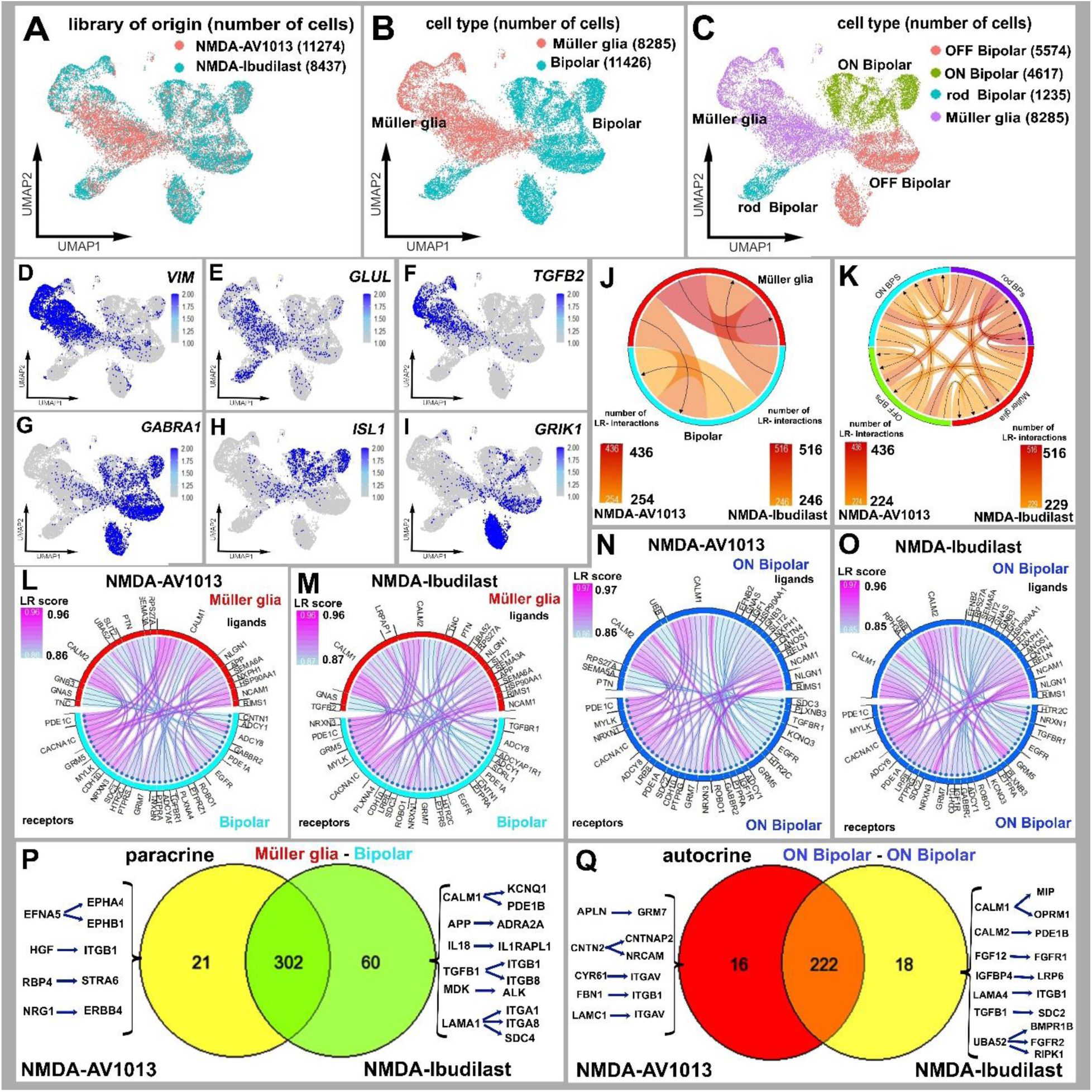
Ligand-receptor interactions were inferred from scRNA-seq data involving MG and BP cells (Bipolar Cells), including ON BP (ON Bipolar Cells) and OFF BP (OFF Bipolar Cells). Retinal cells from libraries treated with NMDA+AV1013 and NMDA+ibudilast were isolated, re-embedded, and organized in UMAP plots of the MG and BP (A-C). MG were identified by the expression of *VIM, GLUL* and *TGFB2* (D, E, F). Bipolar cells were identified by the expression of *GABRA1* (Fig.11 G), ON Bipolar Cells by *ISL1* (H), and OFF Bipolar Cells by *GRIK1* (I). Chord diagrams were used to illustrate potential autocrine and paracrine ligand-receptor (LR) interactions generated from SingleCellSignalR analysis of NMDA+AV1013 and NMDA+Ibudilast libraries (J, K). Chord plots display the 40 most significant paracrine LR interactions from MG to Bipolar Cells (L, M) and autocrine LR interactions in ON Bipolar Cells (N,M) in the NMDA+AV1013 and NMDA+ibudilast libraries. Venn diagram analysis was performed to identify unique and common paracrine and autocrine LR interactions in NMDA+AV1013 vs NMDA+Ibudilast treatments (P, Q). The Venn diagrams illustrate the numbers of unique and common LR interactions between the treatment groups and list representative LR interactions in the paracrine and autocrine conditions of MG and ON bipolar cells.

**Fig 10.**
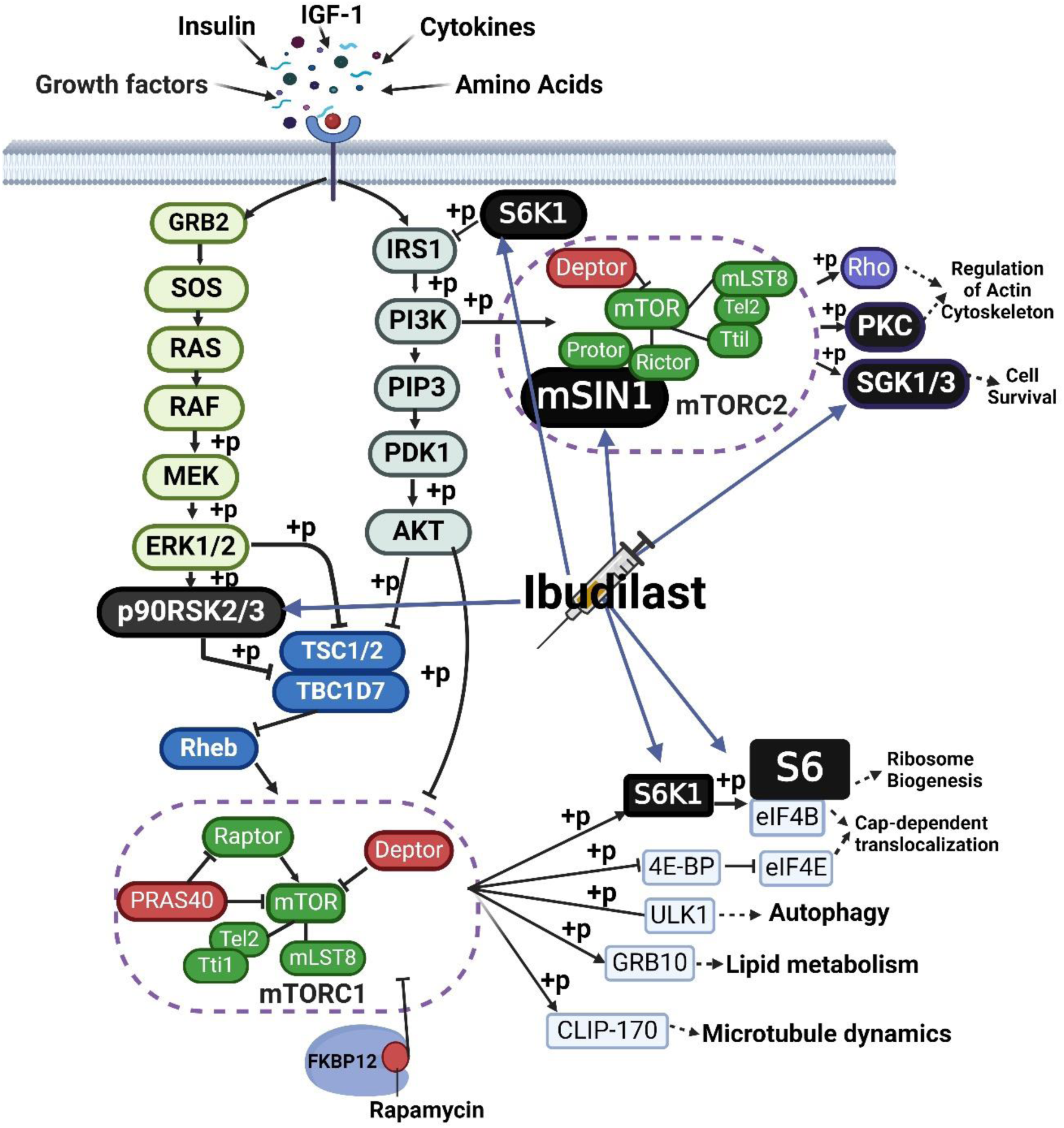
Schematic showing genes upregulated by ibudilast treatment during NMDA retinal damage on scRNA-seq analysis. mTOR signaling pathway diagram, with black background (p90RSK2/3, S6K1, S6, mSIN1, SGK1/3), were identified as upregulated by ibudilast during damage in this study on scRNA-seq analysis. Members of both mTORC1 and mTORC2 were upregulated.

Moreover, the top 40 autocrine LR interactions in ON bipolar cells (Fig 9N, O) were nearly the same between the 2 treatment groups, with only RPH3A-NRXN1 unique to NMDA-ibudilast treatment and PTN-SDC3 unique to NMDA-AV1013 treatment. Although these LR interactions were unique to the top 40, they were present in both ibudilast and AV1013 groups with lower, but still significant, LR scores.

The truly unique LR interactions in the treatment groups were further evaluated with Venn diagram analysis. We identified 383 significant LR interactions from MG signaling to bipolar cells, with 21 unique to only NMDA+AV1013-damaged retina and 60 unique to NMDA+ibudilast-treated retina (Fig 9P). Additionally, we found 256 significant LR autocrine interactions in the ON bipolar group, with 16 unique LR interactions for NMDA+AV1013 and 18 unique LR interactions for NMDA+ibudilast (Fig 9Q).

Within these 2 groups we focused on exploring LR interactions unique to NMDA-ibudilast treatment in both MG and bipolar cells. Both paracrine and autocrine interactions were evaluated. For instance, the unique ibudilast paracrine MG-bipolar cell LR interactions included CALM1-PDE1B/KCNQ1, IL18-IL1RAPL1, MDK-ALK, TGFB1-ITGB1/ITGB8, and LAMA1-ITGA1/ITGA8/SDC4 (Fig 9P). Moreover, the unique ibudilast autocrine LR interactions in ON bipolar cells included CALM1-MIP/OPRM1, CALM2-PDE1B, FGF12-FGFR1, IGFBP4-LRP6, LAMA4-ITGB1, TGFB1-SDC2, and UBA52-BMPR1B/FGFR2/RIPK1 (Fig. 9Q). Notably, many of the unique ibudilast-modulated genes were integrins (e.g., ITGB1, ITGA8) and cytokines/growth factors (e.g., MDK, TGFB1).

## Discussion

In this study we tested the ability of ibudilast to protect the retinal neurons during excitotoxic damage to the eye. In the chick retina, ibudilast, but not its analogue AV1013, decreased NMDA-induced cell death, prevented retinal layer thinning, and preserved the function of the retinal neurons. The fact that ibudilast was protective when given after damage is highly relevant clinically, when medical care may occur hours after the initial injury. Ibudilast preserved different types of neurons in the INL and uniquely protected ON bipolar cell structure and function.

Bipolar cells are critical to vision and play an important role in filtering, processing, and transmitting the light signal from photoreceptors to the RGCs (Nelson & Connaughton, 1995). They synapse with photoreceptors directly, or indirectly via the horizontal cells, and relay the signal to RGCs directly or indirectly via the amacrine cells (Hahn et al., 2023; Masland, 2001). Many subtypes of bipolar cells have been identified in the chick retina by using single cell transcriptomics; as many as 22 distinct types in embryonic retina (Yamagata, Yan, & Sanes, 2021) and 11 distinct types in the retinas of hatched chicks (El-Hodiri et al., 2022)

Structurally bipolar cells are divided into rod and cone bipolar cells based on their synaptic pairing with the corresponding photoreceptor (Bloomfield & Dacheux, 2001; Tsukamoto & Omi, 2017). In mammals, more than 14 cone bipolar cells exist, which represent the diversity of signal processing pathways where bipolar cells are involved. Functionally, however bipolar cells can be categorized into ON and OFF types, depending on their response to the light signal, where ON-center bipolar cells depolarize and OFF-center bipolar cells hyperpolarize in response to increased illumination of their central receptive field (Tsukamoto & Omi, 2017). Islet-1 is described as a marker for ON bipolar cells while PKCa is specific for the rod bipolar cells, which also function as ON bipolar cells (Balaji et al., 2023). Our results showed that ibudilast clearly preserved the ON bipolar cell numbers during NMDA damage, as assessed by Islet-1 staining. Islet-1 expressing horizontal cells were also preserved by ibudilast. In the chick retina ∼50% of the horizontal cells express Islet-1 and TrkA (Fischer, Stanke, Aloisio, Hoy, & Stell, 2007). Moreover, ibudilast preserved ON bipolar cell function as measured by the long flash ON/OFF ERG responses. In contrast, numbers of PKCα positive cell numbers were not different with ibudilast treatment, which suggests that ibudilast specifically protects the cone ON bipolar cells. However, the integrity of the PKCα positive cells was qualitatively better preserved by ibudilast, suggesting an area for future investigation.

The PDE inhibitor component of ibudilast may be critical to its neuroprotective effect, since its analog AV1013, a MIF inhibitor that lacks PDE inhibition, did not share ibudilast’s ability to restore the form and function of the retina. PDE inhibitors are rapidly evolving and increasingly being considered to treat neurologic disorders. In fact, clinical trials are in progress to treat patients with Alzheimer’s disease, autism, depression, and schizophrenia (Delhaye & Bardoni, 2021; Xi et al., 2022). Alternatively, the pharmacodynamics and specificity of ibudilast in the chick retina could favor the protection of inner retinal neurons, whereas AV1013 lacks specificity or is rapidly cleared by cells in the chick retina.

In the retina, PDEs play important and specific roles in conveying the light signal from photoreceptors to the different INL neuronal populations. PDEs hydrolyze cGMP, a second messenger that regulates signaling in nearly every cell class in the retina. cGMP appears to be important in the functioning of ON bipolar cells (Dhingra, Tummala, Lyubarsky, & Vardi, 2014,Nawy, 1990, Shiells, 2002). For example, Shiells et al. found that ON bipolar cell flash responses were potentiated by cGMP (Shiells & Falk, 2002). Our scRNA-seq results confirmed that PDEs affected by ibudilast, especially PDE4B, PDE4D, and PDE10A, are highly expressed in both bipolar cells and MG. Ibudilast’s specific protection of ON bipolar cells might be due to direct inhibition of PDEs in those cells and/or indirectly via inhibition of PDEs in the MG that support and influence bipolar cell function and survival during excitotoxic damage (Bringmann & Wiedemann, 2012). Future studies with conditional knockout of select PDEs in mice could help address the relative importance of cell-specific PDEs.

Moreover, neither drug significantly impacted the morphology of CD45 positive microglia in the retina following damage and had a minimal impact on microglia accumulation. The presence of reactive microglia is known to exacerbate NMDA-induced cell death(Fischer, Zelinka, & Milani-Nejad, 2015b), whereas suppression of microglial reactivity with glucocorticoid agonists suppresses NMDA-induced cell death There were only some weak effects on late accumulation of CD45+ cells in the photoreceptor layer with AV1013. Thus, the neuroprotective effects of ibudilast are likely not mediated by suppressing the reactivity or accumulation of microglia/macrophages.

### mTOR pathway and complex roles in neuroprotection

The mTOR signaling pathway plays a critical role in neuronal survival (Switon, Kotulska, Janusz-Kaminska, Zmorzynska, & Jaworski, 2017). mTOR is a catalytic subunit that interacts with other proteins to form two distinct complexes: mTORC1 and mTORC2, each with unique cellular functions. The mTOR pathway controls many processes that impact major cellular functions like metabolism, protein and lipid synthesis, autophagy, cytoskeletal organization, cellular growth, and survival (Laplante & Sabatini, 2012a, 2012b; Saxton & Sabatini, 2017).

The complex role of mTOR signaling in the retina has been widely documented in the literature. mTOR plays a role in RGC survival and axonal regeneration after optic nerve injury (Wei, Luo, & Chen, 2019). In an optic nerve crush model, conditional knockdown of PTEN and TSC1, negative regulators of mTOR signaling, enhanced RGC survival and regeneration in RGCs positive for mTOR activation, as detected by pS6 staining (Park et al., 2008). The addition of mTOR inhibitor rapamycin, lowered pS6 levels and stalled axonal regeneration in mice with PTEN deletion. This suggests that activation of the mTOR pathway is important in axonal regeneration and survival in the retina. Interestingly, in contrast to neuroprotective findings, others have reported beneficial effects of mTOR pathway inhibition in neurodegenerative disease models (Switon et al., 2017). Ichikawa et al. studied the role of rapamycin in an NMDA induced retinal damage model in rats and found that rapamycin lowered the levels of apoptotic cells and pS6. Swiatkowski et al. found *in vitro* protection of neurons from NMDA damage by inhibiting mTORC1 or GSK3b. Similarly, Ding et al. found that rapamycin plus caspase inhibitor z-vad-fmk could reduce photoreceptor death in experimental retinal detachment, through activation of autophagy. This demonstrates the complexity and context-dependency of mTOR pathway signaling and the importance of tailoring any mTOR pathway intervention to the specific disease setting. While ibudilast upregulated mTOR signaling in the retina during damage, mTORC1 activation was not required for ibudilast’s neuroprotective effect, suggesting that additional pathways upregulated by ibudilast, like HIF1α and Hippo, should be investigated in the future.

### LR/Bipolar cell and MG interaction

The role of MG in maintaining and modulating the function of neurons is well documented (Bringmann et al., 2006). MG interact structurally and functionally with all retinal neurons and consequently contribute to retinal homeostasis and neuroprotection through regulating metabolism, blood flow, ion and water homeostasis, neurotransmitter recycling, and neurotrophic factor release (Bringmann & Wiedemann, 2012). On the other hand, MG can also contribute to neuronal death by accelerating the process of neurodegeneration (Bringmann & Wiedemann, 2012). Our scRNA-seq and ligand-receptor interaction analysis identified many potential autocrine and paracrine interactions of MG and bipolar cells unique to ibudilast treatment in damaged retina.

Our LR interaction analyses suggest some interesting insights on the effect of ibudilast on glia-neuronal interactions in the retina. Although many of the LR pairs identified were similar between the AV1013 and ibudilast treated groups, the unique interactions in the ibudilast group suggest that integrins (e.g., ITGB1, ITGA8) and cytokines/growth factors (e.g., MDK,) may play important roles in ibudilast’s protective function. For example, midkine (MDK), a heparin binding growth factor, has been shown to be neuroprotective when administered exogenously in the chick NMDA retinal damage model (Campbell, Fritsch-Kelleher, et al., 2021). Furthermore, integrins function as cell adhesion receptors involved in cellular communication (Y. G. Mishra & Manavathi, 2021) and may play a role in facilitating neuroprotective signaling from the extracellular matrix to neurons. Santos et al. investigated the involvement of laminin-b1 integrin signaling in retinal ganglion cell survival following retinal ischemia. In a rat retinal ischemia-reperfusion injury model, laminin degradation occurred with decreased b1 integrin activation and loss of RGCs. The RGC death could be inhibited by treatment with agonist antibodies to b1 integrin. We identified LAMA and ITGB1 as unique LR pairs upregulated by ibudilast in the MG and bipolar neurons during NMDA damage and future studies could address the importance of these and other integrin signaling during excitotoxic damage.

Unique to the top 40 paracrine MG->bipolar cells LR interactions in the ibudilast group was LRPAP1 - LRP8/SORL1. LRPAP1 (Low-Density Lipoprotein Receptor-Related Protein Associated Protein 1; also known as receptor associated protein (RAP)) binds to LRP8 receptor and mutations in LRPAP1 are associated with severe myopia and rhegmatogenous retinal detachment (Aldahmesh et al., 2013). It acts as a molecular chaperone that regulates endocytosis and thus influences many other signaling pathways e.g. TGF-β (Ref). LRPAP1 receptor, LRP8, also known as Apolipoprotein E Receptor 2 (ApoER2), plays essential roles in neuronal development and function and its dysregulation may contribute to the pathology of neurodegenerative diseases like Alzheimer’s (Fernandez-Calle et al., 2022). The other LRPAP1 receptor, SORL1 (Sortilin Related Receptor 1), helps regulate endosomal traffic, synaptic protein distribution, and recycling in neurons (Lee et al., 2023; S. Mishra et al., 2022). Further studies are needed to fully understand the role of these receptors in ibudilast protection of retinal bipolar neurons.

## Conclusion

In this study we identified a promising drug, ibudilast, as a protector of retinal form and function during excitotoxic damage in the chick NMDA model. PDE inhibition may be important for ibudilast’s beneficial effect, since its analog AV1013, which lacks PDE inhibition activity, did not protect retinal neurons. Our findings indicate that ON bipolar cells are the main target of ibudilast protection. We utilized scRNA-seq analyses to identity pathways and genes that might be involved in the neuroprotection process. Although ibudilast upregulated mTOR signaling in the retina, mTORC1 signaling was not essential for ibudilast neuroprotection, based on rapamycin inhibitor experiments. We identified several putative LR-interactions between MG and bipolar cells that are unique to ibudilast treatment which might contribute to neuronal preservation. Our findings open the door for future studies that can further investigate mechanisms involved in excitotoxicity and the ability of drugs like ibudilast to overcome damage in common retinal conditions.

## Acknowledgments

This work was supported by the Department of Defense, USA, The Assistant Secretary of Defense for Health Affairs, through U.S. Army Medical Research Acquisition Activity (USAMRAA) under the FY17 Vision Research Program, Technology/Therapeutic Development Award, with Award No.

W81XWH1810805. Additionally, funding was provided by the National Eye Institutes of Health NEI/NIH, USA (R01EY032573), and the Ohio Lions Eye Research Foundation (OLERF). The authors gratefully acknowledge the utilization of the Vision Sciences Research Core Program under P30EY032857 (VSCRP) at The Ohio State University and extend their appreciation to the Cancer Center P30 Core facility of The Ohio State University. The authors also extend their thanks to all ULAR support for animal care at The Ohio State University and special thanks are given to Claire Landreth for ERG assistance and to Mohd Hussain Shah, Richard Wan, Rahaf Shalash, and Misha Sohail for excellent technical assistance.

---

**Figure S1.**
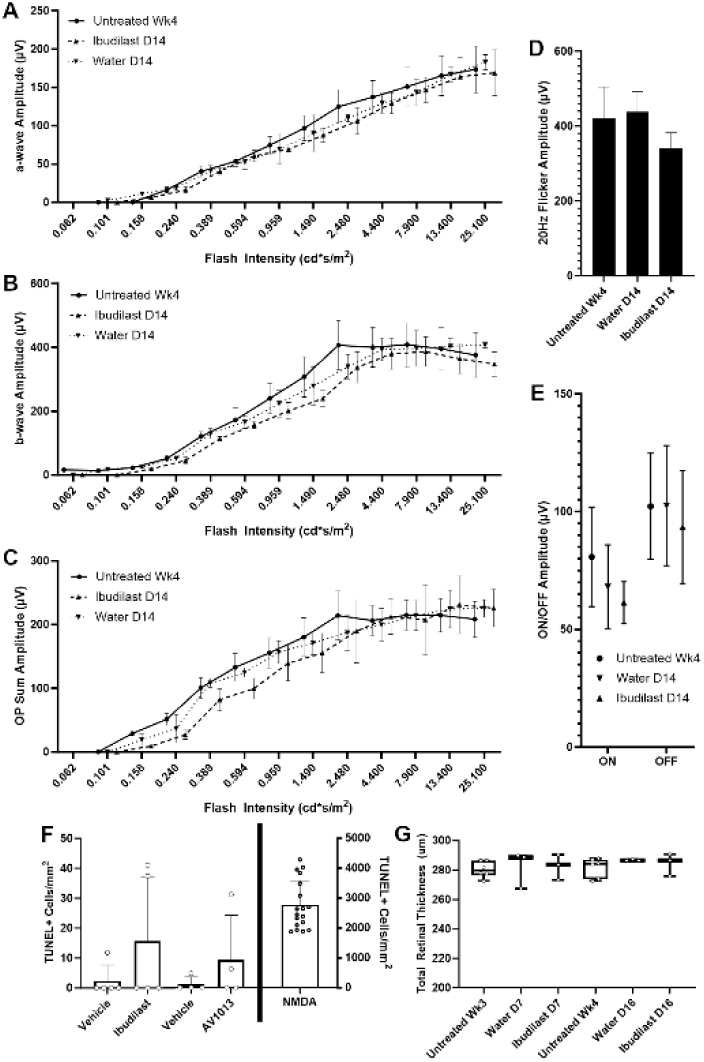
Ibudilast and its analog AV1013 do not induce toxicity in the chick retina. Light adapted ERG shows no difference in waveforms between untreated, ibudilast treated, and vehicle treated undamaged eyes in the following common metrics: a-wave (A), b-wave (B), OP sum (C), flicker (D), and ON/OFF long flash (E). There was no statistical difference in TUNEL positive cell counts 24h after injection of vehicle vs ibudilast or its analog AV1013, while NMDA injection induced large numbers of dying cells at 24h (F). There was no difference in retinal thickness 7-10 days following injection of ibudilast in undamaged eyes, compared to undamaged vehicle or untreated controls (G).

**Figure S2.**
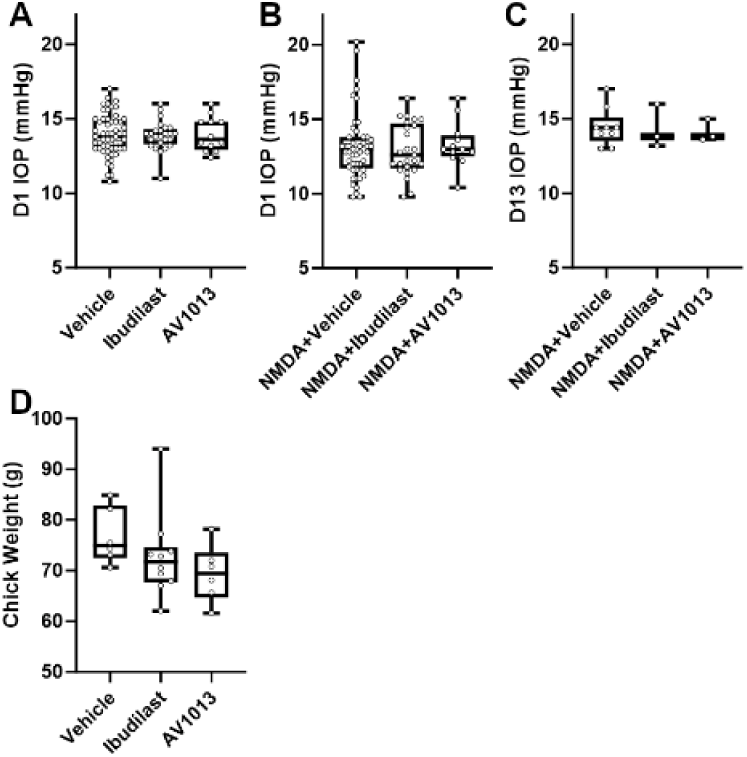
Intravitreal ibudilast and AV1013 does not affect chick intraocular pressure or weight. IOP measurements were acquired in ibudilast, AV1013, and vehicle treated eyes at D1 (A) and in NMDA+ibudilast, NMDA+AV1013, and NMDA+Vehicle treated eyes at D1 (B) and D13 (C). No significant differences were found in either group. No weight differences were found between age-matched chicks treated with ibudilast, AV1013, or vehicle (D).

**Figure S3.**
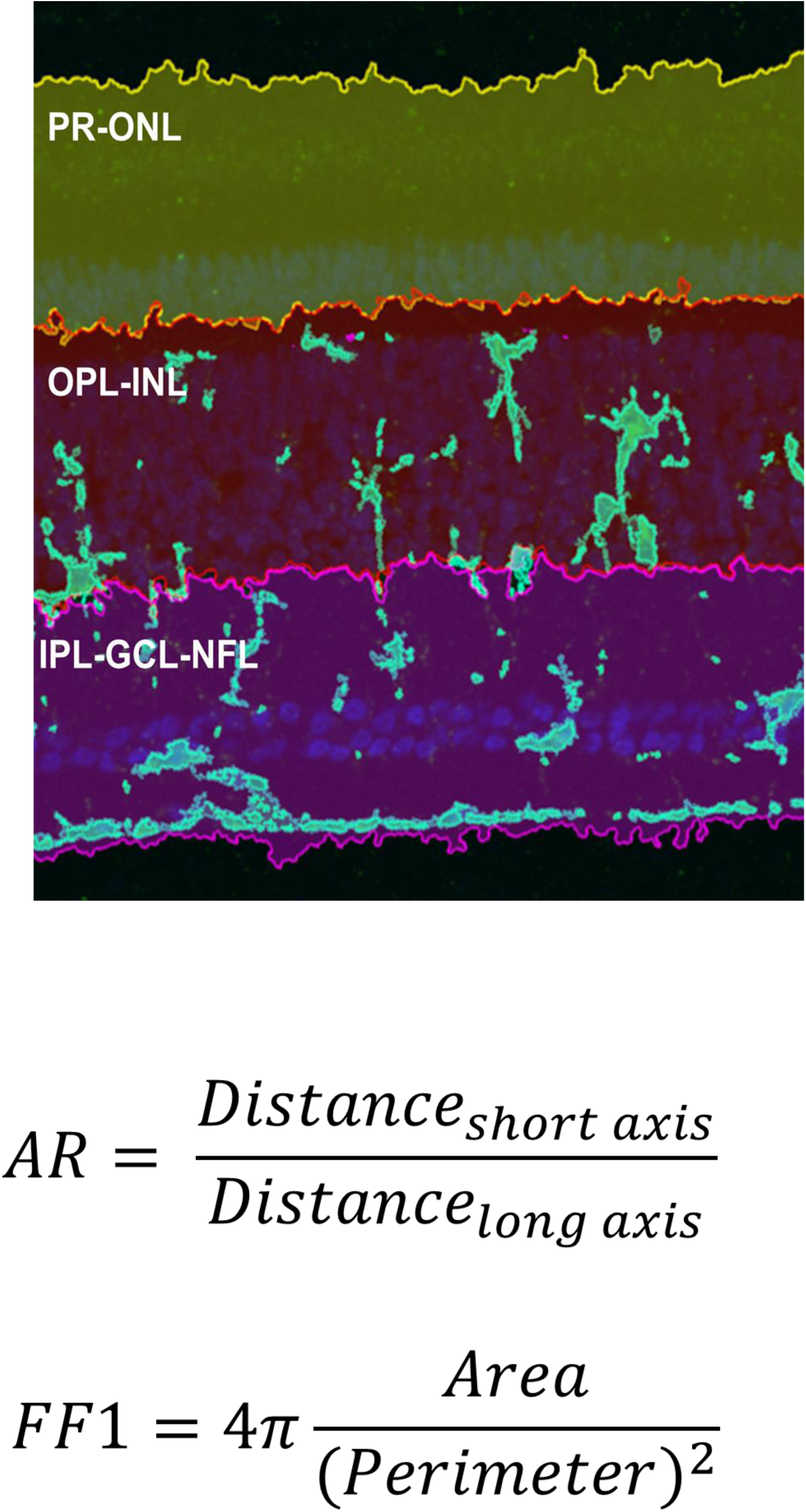
CD45 microglial measurement and morphology analysis by layers. A representative image showing the different regions analyzed for CD45 intensity and morphological parameters. The layers were separated into the PR-ONL, OPL-INL, IPL-GCL-NFL, and the total sum of all three. The aspect ratio (AR) was defined as the short axis over the long axis. Form Factor 1 was defined as 4*π*Area/Perimeter**^2^.**

**Figure S4.**
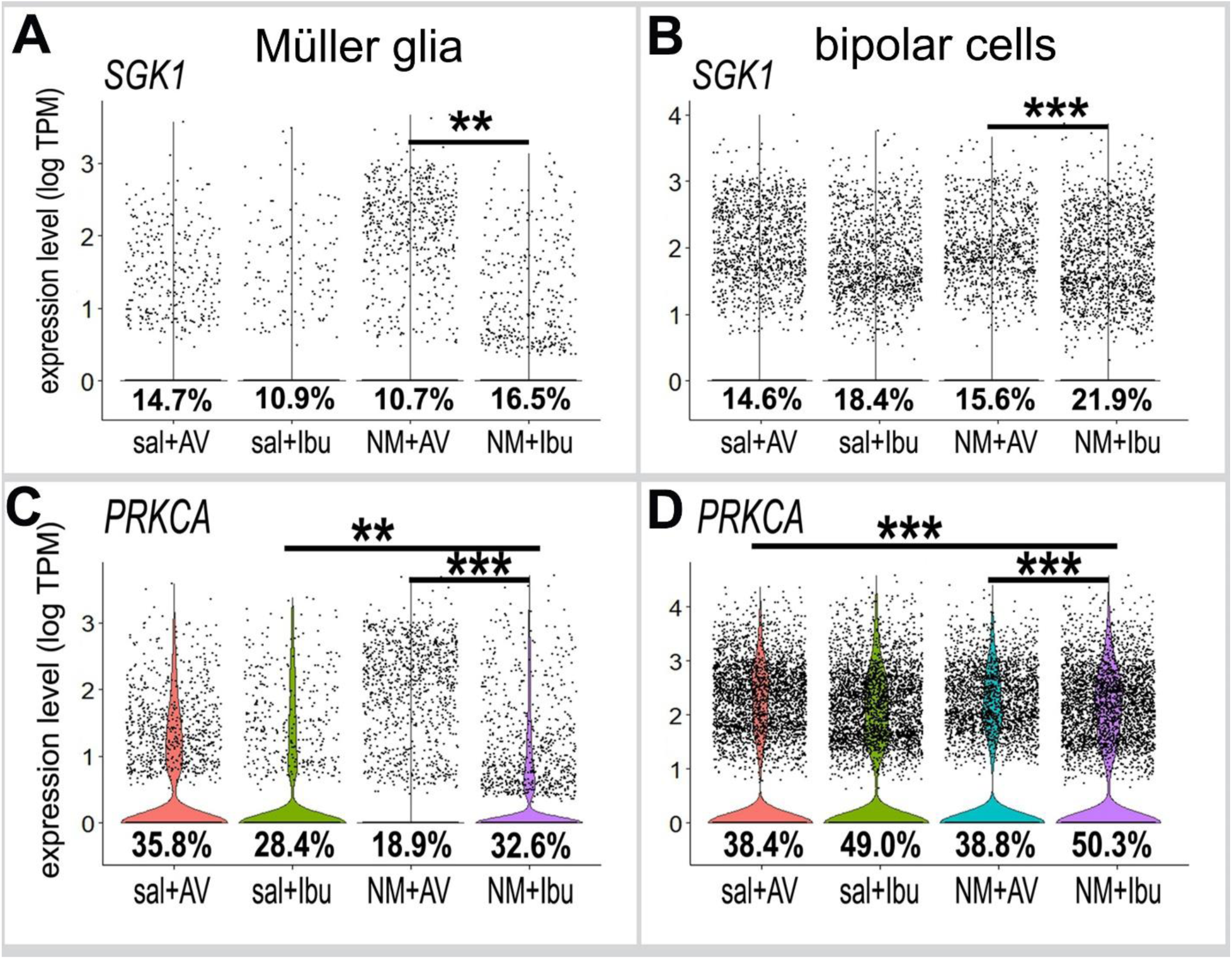
Increased expression of mTOR pathway genes. Violin plots illustrate expression levels of genes related to the mTOR pathway SGK1 (A, B) and PRKCA (C, D) in MG and bipolar cells. **p < 1×10^−5^, ***p < 1×10^−10^ (Wilcoxon Rank Sum Test with Bonferroni correction).

**Table S1.** Statistics for genes in all cells listed in Dot Plot in Figure 1.

|  | **p_val** | **avg_log2FC** | **pct.1** | **pct.2** | **p_val_adj** |
| --- | --- | --- | --- | --- | --- |
| MIF | 0.00E+00 | -0.8604284 | 0.751 | 0.887 | 0.00E+00 |
| CD44 | 0.00E+00 | -0.2858491 | 0.008 | 0.105 | 0.00E+00 |
| PDE10A | 2.44E-36 | 0.3527861 | 0.221 | 0.294 | 5.59E-32 |
| PDE4D | 3.06E-35 | 0.3392327 | 0.576 | 0.616 | 7.00E-31 |
| PDE3A | 5.12E-22 | 0.7466377 | 0.18 | 0.159 | 1.17E-17 |
| PDE4B | 2.99E-14 | 0.5858012 | 0.103 | 0.085 | 6.84E-10 |
| CD74 | 1.09E-01 | 1.1523329 | 0.015 | 0.013 | 1.00E+00 |
| PDE11A | 6.91E-01 | 0.248068 | 0.016 | 0.016 | 1.000000e+00 |

**Table S2.** Statistics for genes in MOiier glia cells listed in Dot Plot in Figure 7.

|  | **p_val** | **avg_log2FC** | **pct.1** | **pct.2** | **p_val_adj** |
| --- | --- | --- | --- | --- | --- |
| EIF4E | 3.65E-84 | -0.1792926 | 0.147 | 0.386 | 8.83E-80 |
| EIF4B | 8.23E-76 | 0.04340044 | 0.135 | 0.353 | 1.99E-71 |
| RPS6KA6 | 1.52E-67 | -0.09096621 | 0.113 | 0.301 | 3.68E-63 |
| MAPKAP1 | 1.92E-55 | -0.07224227 | 0.079 | 0.223 | 4.66E-51 |
| RICTOR | 6.61E-54 | 0.1296343 | 0.14 | 0.325 | 1.60E-49 |
| MAPKAPK2 | 8.70E-51 | -0.01174452 | 0.09 | 0.234 | 2.11E-46 |
| SGK3 | 8.65E-47 | 0.16144606 | 0.149 | 0.325 | 2.10E-42 |
| PDK1 | 2.28E-30 | 0.44240142 | 0.074 | 0.172 | 5.52E-26 |
| PRKCA | 1.69E-20 | 0.40084391 | 0.189 | 0.326 | 4.09E-16 |
| RPS6 | 1.32E-19 | -0.24001174 | 0.601 | 0.783 | 3.20E-15 |
| RPS6KA1 | 3.47E-08 | 0.98731725 | 0.058 | 0.099 | 8.39E-04 |
| SGK1 | 4.99E-08 | 0.72538288 | 0.107 | 0.165 | 1.21E-03 |
| RPS6KA2 | 8.52E-06 | 0.88091899 | 0.193 | 0.161 | 2.06E-01 |

**Table S3.** Statistics for genes in bipolar cells listed in Dot Plot in Figure 7.

|  | **p_val** | **avg_log2FC** | **pct.1** | **pct.2** | **p_val_adj** |
| --- | --- | --- | --- | --- | --- |
| SGK3 | 1.90E-29 | -0.32142832 | 0.269 | 0.38 | 4.60E-25 |
| PRKCA | 1.77E-27 | -0.342425739 | 0.388 | 0.503 | 4.29E-23 |
| RPS6KA1 | 7.07E-16 | -0.211971411 | 0.099 | 0.151 | 1.71E-11 |
| SGK1 | 9.13E-16 | -0.268870669 | 0.156 | 0.219 | 2.21E-11 |
| RPS6KA2 | 4.94E-12 | -0.249947487 | 0.453 | 0.522 | 1.20E-07 |
| MAPKAP1 | 3.24E-08 | -0.124751251 | 0.105 | 0.142 | 7.84E-04 |
| RPS6KA6 | 7.10E-07 | 0.001237986 | 0.092 | 0.124 | 1.72E-02 |
| RICTOR | 9.32E-05 | 0.097351963 | 0.186 | 0.227 | 1.00E+00 |
| EIF4E | 8.64E-04 | 0.240680805 | 0.129 | 0.157 | 1.00E+00 |
| EIF4B | 9.92E-04 | 0.108369536 | 0.106 | 0.13 | 1.00E+00 |
| RPS6 | 3.59E-02 | 0.030735464 | 0.495 | 0.562 | 1.00E+00 |
| MAPKAPK2 | 6.32E-02 | 0.584406426 | 0.019 | 0.024 | 1.00E+00 |
| PDK1 | 6.37E-01 | 0.370922111 | 0.051 | 0.053 | 1.00E+00 |

